# All-optical electrophysiology for high-throughput functional characterization of human iPSC-derived motor neuron model of ALS

**DOI:** 10.1101/289959

**Authors:** Evangelos Kiskinis, Joel M. Kralj, Peng Zou, Eli N. Weinstein, Hongkang Zhang, Konstantinos Tsioras, Ole Wiskow, J. Alberto Ortega, Kevin Eggan, Adam E. Cohen

## Abstract

Human induced pluripotent stem cell (iPSC)-derived neurons are an attractive substrate for modeling disease, yet the heterogeneity of these cultures presents a challenge for functional characterization by manual patch clamp electrophysiology. Here we describe an optimized all-optical electrophysiology, “Optopatch”, pipeline for high-throughput functional characterization of human iPSC-derived neuronal cultures. We demonstrate the method in a human iPSC-derived motor neuron model of ALS. In a comparison of neurons with an ALS-causing mutation (*SOD1* A4V) with their genome-corrected controls, the mutants showed elevated spike rates under weak or no stimulus, and greater likelihood of entering depolarization block under strong optogenetic stimulus. We compared these results to numerical simulations of simple conductance-based neuronal models and to literature results in this and other iPSC-based models of ALS. Our data and simulations suggest that deficits in slowly activating potassium channels may underlie the changes in electrophysiology in the *SOD1 A4V* mutation.

## Introduction

Cell reprogramming technologies have created an unprecedented opportunity to study human neurons *in vitro,* probing disease mechanisms under each patient’s unique genetic constellation (Han et al., 2011, Pankevich et al., 2014). Many studies have used induced pluripotent stem cell (iPSC)-based and direct conversion methods to model neurological, neuropsychiatric, and neurodegenerative diseases, effectively describing disease-related phenotypes in multiple neuronal subtypes (Ichida and Kiskinis, 2015). Here we present methodology for design and analysis of optical electrophysiology experiments on iPSC-based disease models, with application to a model of ALS.

Electrical spiking is the dominant function of every neuron. The spiking patterns, the action potential waveforms, and the sub-threshold voltages under different stimulus waveforms represent an integrative phenotype that reflects the activity of a large number of ion channels, transporters, and pumps, as well as the underlying cellular metabolism. While it is not, in general, possible to deduce the complete ion channel composition of a cell from its spiking patterns (Brookings et al., 2014), differences in spiking patterns between disease-model and control states can point to likely differences in ion channel function; and pharmacological rescue of disease-associated functional phenotypes can support efficacy of a candidate therapeutic.

Electrophysiology data has been traditionally difficult to attain. Manual patch clamp measurements can be highly accurate, but are labor-intensive and slow. Multi-electrode arrays and calcium imaging probe overall spontaneous activity of a culture, but do not probe details of action potential waveforms, nor are these techniques typically combined with precisely targeted stimulation. The large effort required to record manually from many neurons, combined with the intrinsic variability of iPSC-derived cultures, presents a major obstacle to systematic exploration of patient populations or experimental conditions.

A recently developed system for all-optical electrophysiology (“Optopatch”) addresses this bottleneck (Hochbaum et al., 2014). Optogenetic actuation occurs through a blue light-activated channelrhodopsin, called CheRiff. Voltage imaging occurs through a spectrally orthogonal near infrared genetically encoded fluorescent voltage indicator, called QuasAr2. Specialized optics and software allow simultaneous stimulation and recording from multiple single cells embedded in a complex network (Werley et al., 2017, Zhang and Cohen, 2017). However, low expression levels of the Optopatch construct limited application to highly robust primary neuron cultures and to commercially produced iPSC iCell neurons (Hochbaum et al., 2014). Furthermore, limitations in data handling and analysis constrained previous applications to relatively small numbers of well separated neurons.

Scaling up the Optopatch platform for iPSC-based disease modeling posed a number of challenges in automated data analysis and statistical interpretation. We developed image segmentation techniques to extract the fluorescence traces and morphology of individual neurons, even when they were clumped and overlapping. We developed a novel suite of filtering and fitting techniques robust to the dominant noise sources in our dataset to extract spike times and action potential waveform parameters (Cunningham and Byron, 2014, Druckmann et al., 2013). We then employed systematic regression techniques to determine population- and subpopulation-level differences between the mutant and control cell lines while controlling for significant sources of cell-to-cell variability.

Here we apply Optopatch assays to study the electrical properties of human iPSC-MNs in a model of ALS. We developed improved expression constructs and cell culture protocols to measure spontaneous and optogenetically induced spiking in human iPSC-derived MNs. We applied these tools to a previously validated model of ALS with the *SOD1* A4V mutation, and its gene-corrected but otherwise isogenic control. We measured 1771 single cells across 6 differentiations, for mutant and control, in two independent isogenic pairs. We found that *SOD1 A4V* mutant cells had higher spontaneous activity than isogenic controls, and greater firing rate at low stimulation, but lower firing rate under strong stimulation due to an increased likelihood of entering depolarization block. Mutant cells also had smaller amplitude action potentials. Mutant and genome-corrected cells had indistinguishable maximum firing rates and intra-stimulus accommodation behavior.

To gain mechanistic insight into this array of seemingly distinct functional comparisons, we explored simplified conductance-based Hodgkin-Huxley-type models. Variation of a delayed rectifier potassium channel was sufficient to account for the bulk of our findings. The relative ease of acquiring Optopatch data creates an opportunity to explore electrophysiology in cell-based models of neurological disease in detail and at a population scale and to make quantitative comparisons to theory.

## Results

### Expression and characterization of Optopatch in human iPSC-derived motor neurons

We developed an experimental pipeline to apply Optopatch to an established (Kiskinis et al., 2014, Wainger et al., 2014) human iPSC-based model of ALS (Fig. 1A). The major steps were: (1) differentiation of iPSCs into MNs, (2) delivery of Optopatch genes, (3) optical stimulation and recording, (4) image segmentation, (5) voltage trace parameterization, (6) statistical analysis of population differences, and (7) comparison to numerical simulations. We applied the pipeline to two iPSC lines: one derived from an ALS patient (39b) harboring the A4V mutation in the *SOD1* gene, the other an isogenic control cell line (39b-Cor), generated by correcting the mutation in *SOD1* through zinc finger nuclease (ZFN)-mediated gene editing. Both lines have been extensively characterized and validated for pluripotency markers, developmental potency, and genomic integrity as described previously(Kiskinis et al., 2014, Wainger et al., 2014). We validated the key results in a second patient-derived line with the same mutation in *SOD1* (RB9d), and a corresponding isogenic control line (RB9d-corr) (Figure S1A,B).

**Figure 1.**
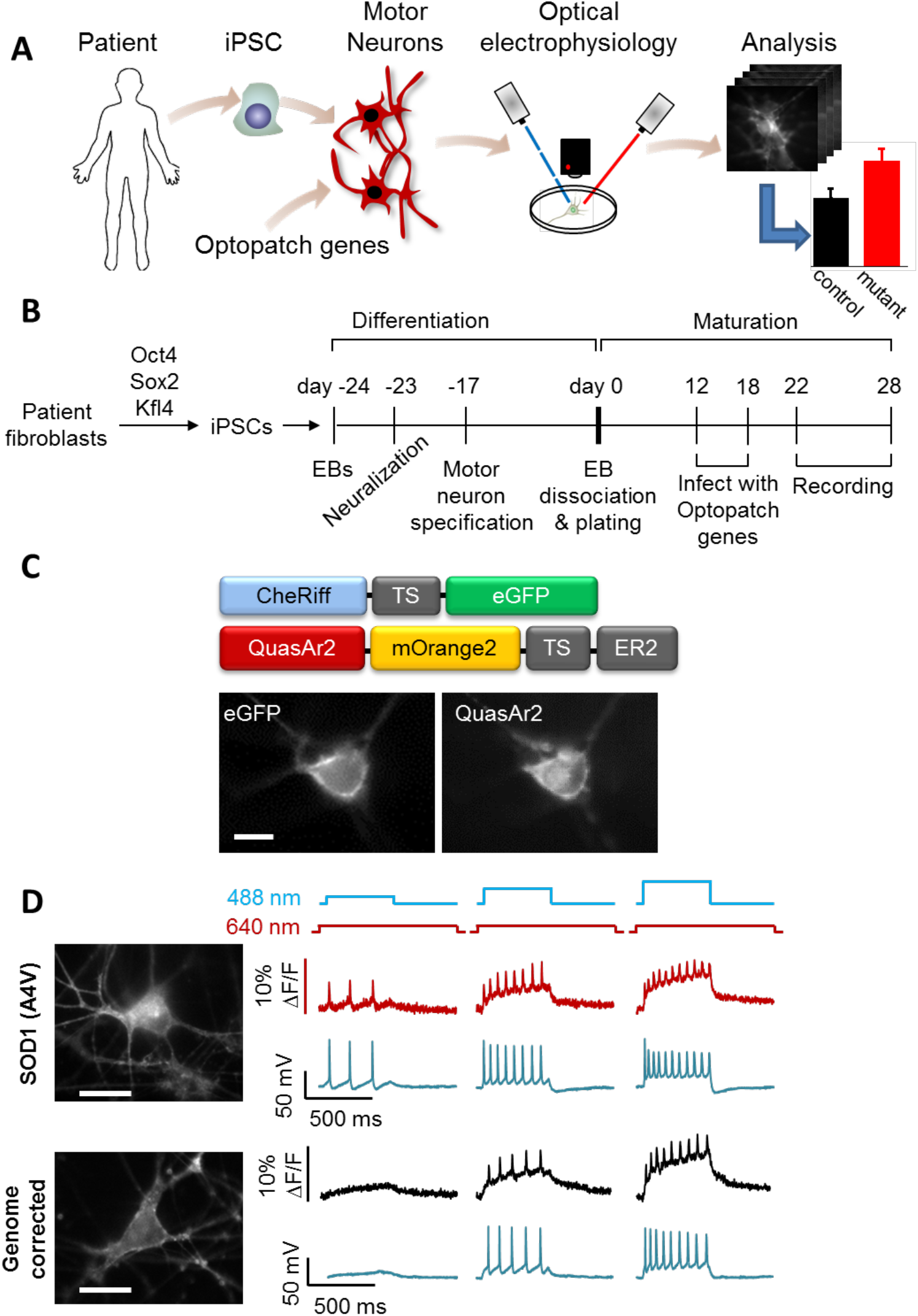
Optopatch reports firing patterns of iPSC-derived motor neurons in a model of ALS. **(A)** Pipeline for disease modeling with optical electrophysiology. **(B)** Timeline of motor neuron differentiation, gene transduction, maturation, and measurement. **(C)** Top: domain structure of Optopatch constructs. Bottom: images of an iPSC-derived motor neuron expressing both CheRiff-eGFP and QuasAr2-mOrange2. **(D)** Simultaneous fluorescence and patch clamp recordings of spiking in iPSC-derived motor neurons under optical stimulation. Left: Images from mutant and genome corrected controls. Right: Fluorescence (red, black) and voltage (blue). Illumination protocol shown above. Scale bars in all panels 10 μm. See also Supplemental Fig. S1.

We differentiated the iPSC lines into postmitotic, spinal MNs using a previously described protocol based on formation of embryoid bodies and subsequent neuralization through dual-SMAD inhibition (Fig. 1B). MN specification was achieved through addition of retinoic acid and a smoothened agonist (Kiskinis et al., 2014, Boulting et al., 2011). We and others have previously shown that the majority of MNs generated through this protocol are FOXP1/HOXA5 positive, indicative of a lateral motor column identity with a rostral phenotype, and are able to form neuromuscular junctions (Kiskinis et al., 2014, Amoroso et al., 2013). This 24-day protocol resulted in highly neuralized cultures (> 95% MAP2/TUJ1^+^ cells) and significant numbers of spinal MNs (> 30% of MAP2/TUJ1^+^ were ISL1/2 (ISL)^+^) (Fig. S1A, B). At the end of the differentiation, MN cultures were plated onto poly-D-lysine/laminin-coated glass-bottom dishes for subsequent maturation and electrophysiological analysis.

We tested the calcium-calmodulin dependent kinase II type-α (CamKIIα) promoter as a means to achieve selective and specific expression in iPSC-MNs. Previously published RNA-Seq data acquired from FACS-isolated HB9^+^ MNs differentiated through this protocol (Kiskinis et al., 2014) revealed strong expression of CAMK2A (Fig. S1C). The CaMKIIα promoter is known to be active in mature excitatory neurons (Lund and McQuarrie, 1997). To validate the specificity of the CamKIIα promoter for MNs we infected iPSC-derived MN cultures with a CamKIIα-eGFP lentiviral construct and performed immunocytochemistry for eGFP and ISL (Fig. S1D). Of the ISL^+^ MNs, 75% expressed eGFP. Of the eGFP^+^ cells, 89% were also ISL^+^ MNs (*n* = 1147 ISL^+^ MNs, 1289 eGFP^+^ cells; Fig. S1E).

The previously published Optopatch construct (Hochbaum et al., 2014) contained the CheRiff and QuasAr2 genes joined by a self-cleaving 2A peptide. We found that this construct did not express highly enough for robust functional recordings in iPSC-MNs. The expression level was considerably higher when the two genes were packaged in separate lentiviruses. We generated low-titer lentiviruses (Methods) for QuasAr2-mOrange2 (Addgene #51692) and CheRiff-eGFP (Addgene #51693), both under control of the CamKIIα promoter (Fig. 1C). We delivered the Optopatch genes via lentiviral transduction of the MN cultures 10 days before each recording.

Neurons were imaged 22-28 days post-plating (DPP). Neurons showed robust expression of eGFP, indicative of CheRiff expression, as well as near infrared fluorescence indicative of QuasAr2 expression (Fig. 1C). Both proteins showed extensive membrane trafficking in the soma and in distal processes, though QuasAr2 also showed some intracellular puncta.

Optopatch expression was reported not to have significant effects on electrical properties of primary or iPSC-derived neurons (Hochbaum et al., 2014) but those measurements did not include MNs. To test for effects of expression on MN electrophysiology, we performed manual patch clamp measurements in iPSC-MNs expressing either CaMKIIα-driven Optopatch or a CaMKIIα-driven eGFP control (Methods). Optopatch expression did not significantly perturb resting voltage, membrane resistance, membrane capacitance, rheobase, or AP threshold voltage relative to control cultures expressing eGFP (Fig. S2A-E). Both Optopatch and eGFP-transduced cells had slightly lower membrane resistance and higher membrane capacitance than non-transduced controls, indicating that within the heterogeneous *in vitro* neuronal population, the CaMKIIα promoter targeted expression to neurons with larger surface area, a marker of greater maturity. We further tested for differences in action potential properties between eGFP- and Optopatch-expressing MNs, in both *SOD1 A4V* and genome corrected controls. Individual cells showed widely varying firing patterns, particularly in the vicinity of depolarization block, ranging from single spikes to tonic firing. Many cells showed subthreshold ringing oscillations that gradually diminished as the cell entered depolarization block. In neither genotype did we observe significant differences between eGFP- and Optopatch-expressing MNs in action potential amplitude, maximum firing rate, or width of the first spike following stimulus onset (Fig. S2F-H).

To test whether the optical measurements were a faithful reporter of action potential waveforms, we acquired simultaneous manual patch clamp and optical recordings, with optogenetic stimulation (Methods; Fig. 1D). Fluorescence traces were extracted from single-cell recordings using a previously described pixel weighting algorithm which automatically identified pixels whose fluorescence correlated with the whole-image mean (Kralj et al., 2012). Action potentials (APs) were resolved optically on a single-trial basis and tagged with an automatic spike-finding algorithm (Supplemental Methods). Of the optically identified spikes, 4% were not automatically identified in the patch clamp recordings; of the electrically identified spikes, 5% were not automatically identified in the optical recordings (*n* = 148 spikes, 4 cells). These discrepancies came not from shot noise (which contributed an error rate < 10^-6^) but from low-amplitude oscillations near depolarization block whose classification as spike or not-spike was ambiguous even to human observers (see Fig. 6B for an example).

We compared AP parameters of spikes recorded simultaneously optically and electrically. Optically recorded APs had a root-mean-square (r.m.s.) error of 1.2 ms in time of peak depolarization relative to the electrical signal (*n* = 148 spikes, 4 cells), and a systematic overestimate of spike full-width at half-maximum (FWHM) by 1.8 ± 1.1 ms (mean ± s.d.) compared to an electrically recorded mean spike FWHM of 5.6 ms (Fig. S2I). These mean widths included exceptionally broad spikes near onset of depolarization block. The average width for the first electrically recorded AP after stimulus onset was 3.9 ms, comparable to the literature on developing motor neurons (in rat, AP duration is 9.3 ms at E15-16 and 3.4 ms at P1-3, at 27-29 °C (Ziskind-Conhaim, 1988)). In the electrical recordings, the first AP after stimulus onset was narrower in width (3.9 ms vs. 6.7 ms FWHM) and higher in amplitude (74 mV vs. 60 mV), than subsequent APs. In the optical recordings the first AP was also narrower than subsequent APs (5.7 ms vs 8.7 ms), but appeared smaller in amplitude (2.5% ΔF/F vs. 2.8% ΔF/F) than subsequent APs. The differing trends in apparent spike amplitude are explained by low-pass filtering of the optical signal due to the 2 ms exposure time of the camera and the 1.2 ms response time of QuasAr2. This level of time resolution enabled robust spike counting and coarse parameterization of AP waveforms but not detailed analysis of sub-millisecond dynamics.

The fluorescence signal showed a slow increase in baseline during each optical stimulus epoch (Fig. 1D), which we traced to blue light photo-production of a red-fluorescent product, as has been reported previously for other Arch-derived voltage indicators (Venkatachalam et al., 2014, Hou et al., 2014) (Supplemental Discussion). This effect had been negligible in previous experiments in primary rodent neurons (Hochbaum et al., 2014) on account of higher CheRiff expression (necessitating lower blue stimulus intensity) in the primary cells. We did not include gradual changes in baseline in the analysis, focusing instead on spike timing and shape parameters.

### Probing neuronal excitability with Optopatch

Fig. 2A shows the illumination protocol used to probe the cell-autonomous excitability of human iPSC-MNs. Recordings were acquired at 500 Hz frame rate, for 9,000 frames. Illumination with red light (635 nm, 800 W/cm^2^) induced near infrared voltage-dependent fluorescence of QuasAr2. Cells were monitored for 10 s without stimulation to quantify spontaneous activity. Cells were then stimulated with eight 500 ms pulses of blue light of linearly increasing intensity from 6 mW/cm^2^ to 100 mW/cm^2^. After each blue stimulus pulse, cells were recorded for another 500 ms without blue stimulus, and then given 5 s of rest with neither red nor blue illumination. Mutant *SOD1* A4V and genome corrected control MN cultures were differentiated in parallel and recordings from paired cultures were performed on the same day.

**Figure 2.**
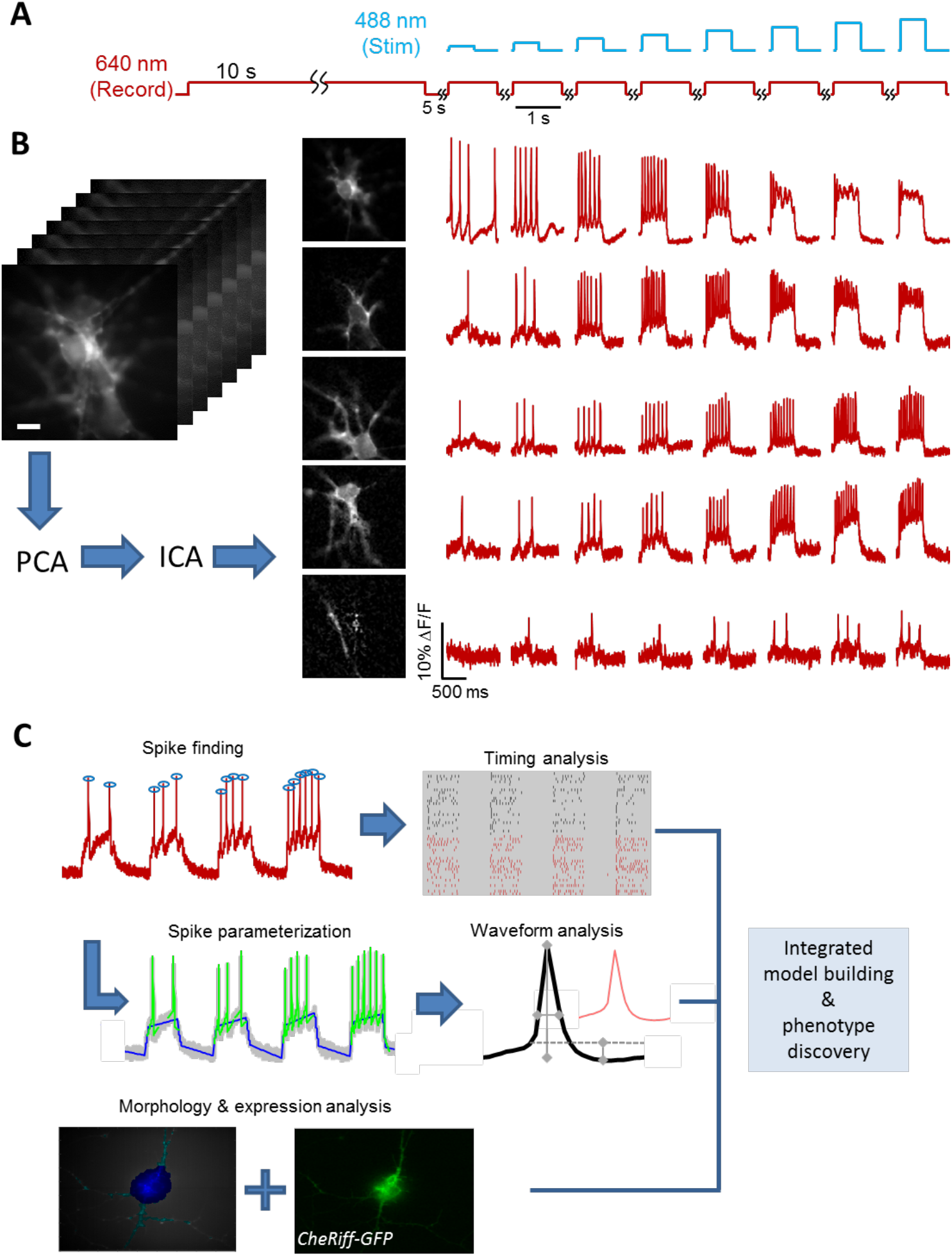
Optopatch measurement and analysis pipeline. **(A)** Cells were subject to 10 s of unstimulated recording to measure spontaneous activity (red), and then to 8 stimulation pulses of 500 ms duration and increasing intensity (blue). **(B)** Activity-based movie segmentation. Image stacks were filtered spatially and temporally, and then processed via principal components analysis (PCA) followed by independent components analysis (ICA) to identify clusters of pixels whose fluorescence values co-varied in synchrony. The movie was decomposed into a sum of overlapping neuron images, each with its own spiking pattern. Scale bar 10 μm. **(C)** Components of the parameterization pipeline. Spikes were identified in the fluorescence traces. Spiking patterns were analyzed within stimuli, between stimuli and between populations. Action potential waveforms were also parameterized, enabling comparison of width, height, and after-polarization within and between cells. The results of the segmentation in the spatial domain enabled measurement of morphological features (soma vs dendrite) and of CheRiff-eGFP expression level. Finally, all of this information was integrated to build a coherent picture of phenotypic differences between mutant and control cell lines. See also Supplemental Figs. S2 – S4.

### Image segmentation and data processing

To accommodate the large quantities of Optopatch data (1039 movies, 200 gigabytes total), we developed a pipeline for analysis in a parallel computing environment (Fig. S3). The first stage comprised image segmentation. Cells often clustered, with overlapping somas and intertwined processes. On average, 60% of each cell body area overlapped with other cells in the controls, and 63% in the mutants (distribution difference *p* = 0.07 by Mann-Whitney U test, not significant). The large degree of overlap implied a need to un-mix the fluorescence signals from overlapping cells.

Our segmentation approach was derived from an independent components analysis (ICA) algorithm, originally developed for calcium imaging (Mukamel et al., 2009), with modifications to accommodate the differing morphological, statistical and noise properties of voltage imaging data (Supplemental Methods). In brief, movies were high-pass filtered in time to accentuate the signals from spikes, and low-pass filtered in space to suppress spatially uncorrelated shot noise. Movies were then subjected to principal components analysis (PCA) to reduce the dimensionality of the dataset, and then time-domain ICA to identify linear combinations of principal components that maximized statistical independence between intensity traces. The spatial filters produced from ICA were then applied to the original (unfiltered) movie to extract the underlying intensity traces (Supplemental Methods; Fig. 2B).

Complex images of up to six overlapping neurons were readily decomposed into single-cell traces. No information about cell morphology was used in the image segmentation, so emergence of neuron-shaped objects with corresponding stereotyped AP waveforms confirms the effectiveness of the algorithm. Inspection of the firing patterns revealed negligible crosstalk between signals derived from overlapping cells. Single-cell fluorescence traces were then processed with a spike-finding algorithm that used a dynamically adjusted threshold to accommodate different signal-to-noise ratios in different cells (Supplemental Methods; Fig. 2C). Sources were classified as active cells if they showed five or more spikes during the experiment and had a signal to noise ratio greater than 5, ensuring a shot noise contribution to errors in spike calling of < 10^-6^.

The second stage of the pipeline comprised parameterization of the spike waveforms. A standard set of parameters has been proposed to describe AP waveforms recorded via conventional electrophysiology (Druckmann et al., 2013). Fluorescence differs from patch clamp in that fluorescence has a lower signal-to-noise ratio, does not have an absolute voltage scale, is subject to baseline drift, and has lower time resolution. To determine what parameters we could use robustly, we first described spikes with a large set of parameters and then used an information-theoretic approach to eliminate redundancies (Supplemental Discussion; Fig. S4). Our final set of parameters described the upstroke duration, downstroke duration, initiation threshold relative to baseline, spike amplitude relative to baseline, and after-hyperpolarization relative to baseline.

Our image segmentation method also enabled quantitative description of cellular morphology. We used filters derived from the activity-based segmentation to identify the two-dimensional footprint of each cell. We then employed morphological image processing to identify cell soma and dendrites (Supplemental Methods; Fig. S4).

### Comparison of mutant *SOD1* A4V vs isogenic control MNs

We compared the firing patterns between iPSC-MNs with the *SOD1* (A4V) mutation (n = 331) and genome-corrected controls (n = 843). Fig. 3A shows a raster plot of the spike timing for a subset of the cells (not including the recording of spontaneous activity at the start of each trace). Fig. 3B shows that, on average, *SOD1 A4V* cells had higher spontaneous activity than genome corrected controls (mean spontaneous rates 1.50 ± 0.18 Hz in mutant, 0.98 ± 0.12 Hz in control, *p* = 0.003, Wilcoxon signed rank test used because of non-normal distribution). Fig. 3C shows the population-average spike count in a 500 ms stimulus as a function of stimulus intensity for the two genotypes. The curves for mutant and control crossed: in the most strongly stimulated epoch, mutant cells fired less on average than controls (mean rates 12.6 ± 0.5 Hz in mutant, 14.6 ± 0.3 Hz in control *p* = 0.0012, unpaired *t-test*). We then performed matched experiments in a second patient-derived *SOD1* (A4V) cell line (RB9d) and its isogenic control (RB9d corr). As with the 39b line, the RB9d mutant cells showed, on average, enhanced spontaneous activity, hyperexcitability at weak stimulus, and hypoexcitability at strong stimulus (Figure S5).

**Figure 3.**
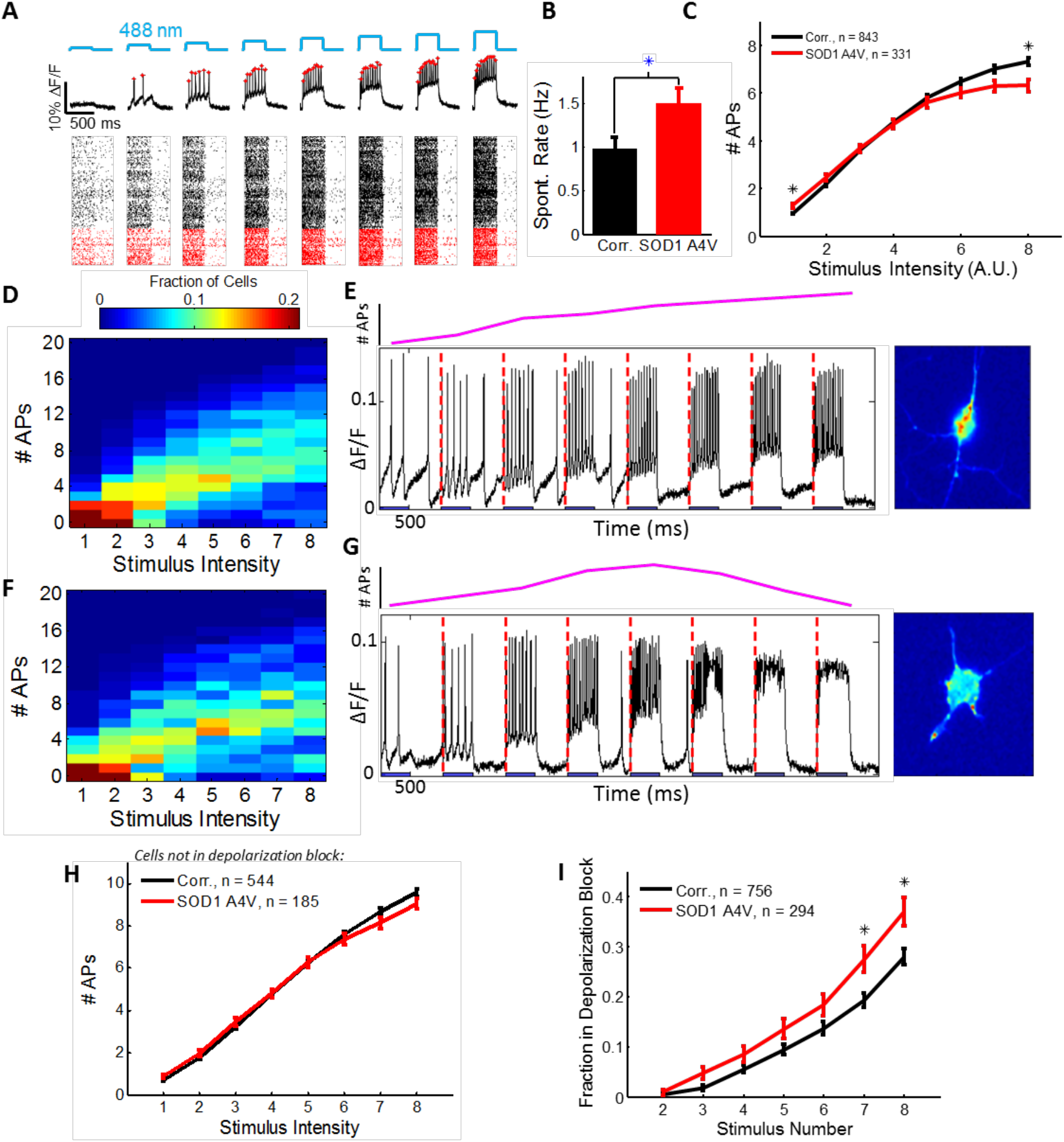
Comparison of spiking patterns in *SOD1* A4V and control motor neurons. **(A)** Top: representative optical trace with stimulus protocol (blue) and identified spikes (red stars). Bottom: Raster plot showing spike timing for a subset of the recorded cells. Black are controls, red are *SOD1* A4V. **(B)** Spontaneous activity. **(C)** Population-average number of action potentials as a function of optogenetic stimulus strength. **(D, F)** Histograms of number of APs as a function of stimulus intensity for (D) controls and (F) *SOD1* A4V mutants. **(E)** Left: Spike train in a control neuron showing monotonically increasing number of APs as a function of stimulus strength. Right: Image of the cell. **(G)** Left: Spike train in a *SOD1* A4V mutant showing depolarization block upon strong stimulus. Right: Image of the cell. **(H)** Number of APs as a function of stimulus strength among the sub-population of cells that did not enter depolarization block. No significant differences were observed between mutant and control. **(I)** Fraction of cells in depolarization block as a function of stimulus number. Stars indicate significance differences between mutant and control to *p* < 0.01. See also Supplemental Figs. S5, S6.

Population-level differences in activity could arise from uniform shifts in all cells or from redistribution of cells among sub-populations with different firing patterns. The single-cell Optopatch data allowed us to examine the underlying distributions of single-cell behavior that led to the population-average differences (Fig. 3D for controls, 3F for mutants).

Under strong stimulus, two clear sub-populations emerged: cells that fired rapidly and tonically (Fig. 3E), and cells that generated just one or two spikes before going quiet (Fig. 3G). We presumed that these quiet cells were constitutively inactive. However, a majority of these cells (64% in the control, 72% in the mutant) fired four or more times during a stimulus of intermediate intensity. These results established that a sub-population of neurons showed a non-monotonic dependence of firing rate on stimulus strength, with a maximum in firing rate at intermediate stimulus strength.

We then examined the fluorescence waveforms of the cells that inactivated under strong stimulus. These cells were depolarized but not firing, a signature of depolarization block (Pontiggia et al., 1993). To quantify the populations in depolarization block, we defined the onset of depolarization block as a decrease in number of spikes upon an increase in stimulus strength. Cells that did not show depolarization block had statistically indistinguishable firing rates in mutant and control populations at high intensity (*p* = 0.11 unpaired t-test at the strongest stimulus; Fig. 3H). However, the proportion of cells that entered depolarization block differed significantly between mutant and control: at the strongest stimulus, the *SOD1* A4V cells were 32% more likely to be in depolarization block than the controls (*p* = 0.008 binomial model t-test with Holm-Bonferroni correction, two hypotheses; Fig. 3I). This difference in propensity to enter depolarization block under strong stimulus was the most dramatic difference between mutant and control.

Aligning all cells’ F-I curves by the stimulus pulse at which they reached their maximum firing rate revealed a stereotyped F-I curve (Fig. 4). Cells showed linear dependence of firing rate on stimulus strength up to a maximum, maintenance of the maximum firing rate in a plateau phase, and then a rapid collapse into depolarization block. The mutant population tended to have a narrower plateau phase and a greater propensity to enter depolarization block than the controls. For cells with matched firing rates at a given stimulus strength, the odds of a cell going into depolarization block in the next stronger stimulus were 88% higher in the mutants than the controls (p = 1.7 × 10^-4^ logistic regression coefficient t-test). The mutant and control neurons reached the same maximum firing rate (mean rates 8.7 ± 0.4 Hz in mutant, 9.2 ± 0.3 Hz in control *p* = 0.51, linear model coefficient t-test).

**Figure 4.**
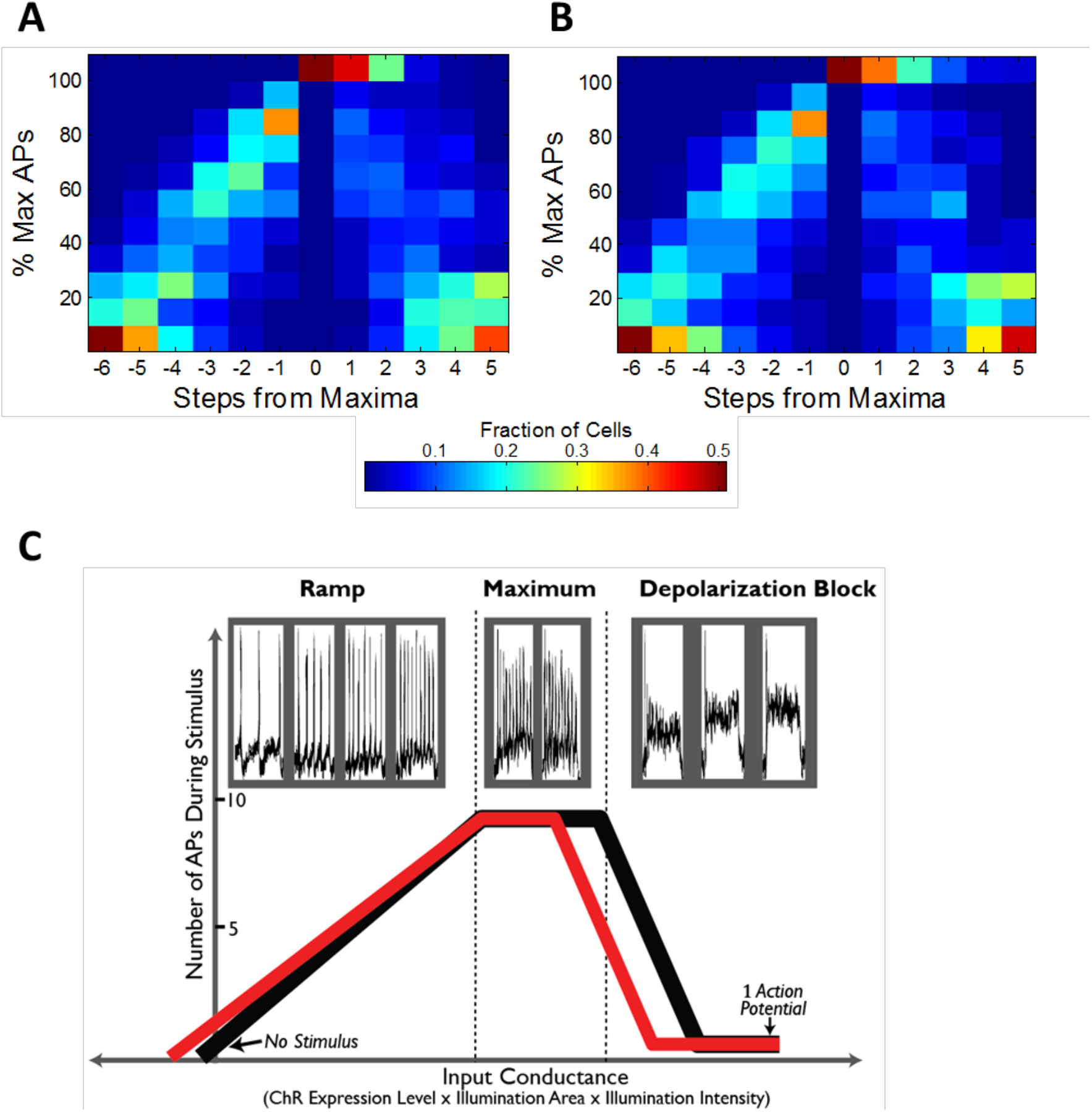
Characteristic firing pattern of iPSC-MNs as a function of stimulus strength. **(A)** Heat maps showing universal shape of the F-I curves for control and *SOD1* (A4V) mutant iPSC-MNs. F-I curves from each cell were rescaled along the x- and y-axes as follows. Firing rate was expressed as a percentage of the cell’s maximum firing rate. Stimulus intensities were aligned to the lowest intensity stimulus at which this maximum firing rate was achieved. This rescaling revealed typical F-I trajectory shapes in a manner that was independent of changes in CheRiff expression level. **(A)** Control and **(B)** *SOD1 A4V* mutant iPSC-MNs showed a linear increase in firing rate vs. stimulus strength for weak stimuli, a plateau in firing rate for moderate stimuli, and a collapse in firing under strong stimuli. **(C)** Top: Fluorescence traces from a single representative cell which passed through three distinct stages of firing in response to monotonically increasing optogenetic stimulus strength. Bottom: Fit of piecewise-continuous F-I curves to the data in (A) and (B) for (black) genome corrected and (red) *SOD1* A4V mutant cell lines. Curves were constructed from measurements of spontaneous rate, average slope of the F-I curve (controlling for expression level), maximum firing rate, and the number of stimulus steps spent at the maximum.

We next studied the timing of the APs within each stimulus epoch. After the first two APs, cells showed nearly constant firing rate throughout the stimulus, provided that the cell was not in depolarization block (Fig. 5). To quantify the degree of firing rate adaptation, we defined the *n*^th^ inter-spike interval by *ISI*_n_ = *t_n+1_* – *t*_n_, where *t*_j_ represents the time of peak of spike *j*. We then defined the degree of firing rate adaptation by: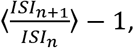 where the average is over all spikes during a stimulus epoch. This quantity did not differ significantly between *SOD1* A4V and control neurons (8.7% in control, 8.9% in mutants, p = 0.87 unpaired t-test). We also detected no differences between mutant and control in the ratio of first inter-spike interval to the average inter-spike interval during a stimulus epoch 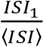 (33% in control, 34% in mutant, p = 0.45 unpaired t-test).

**Figure 5.**
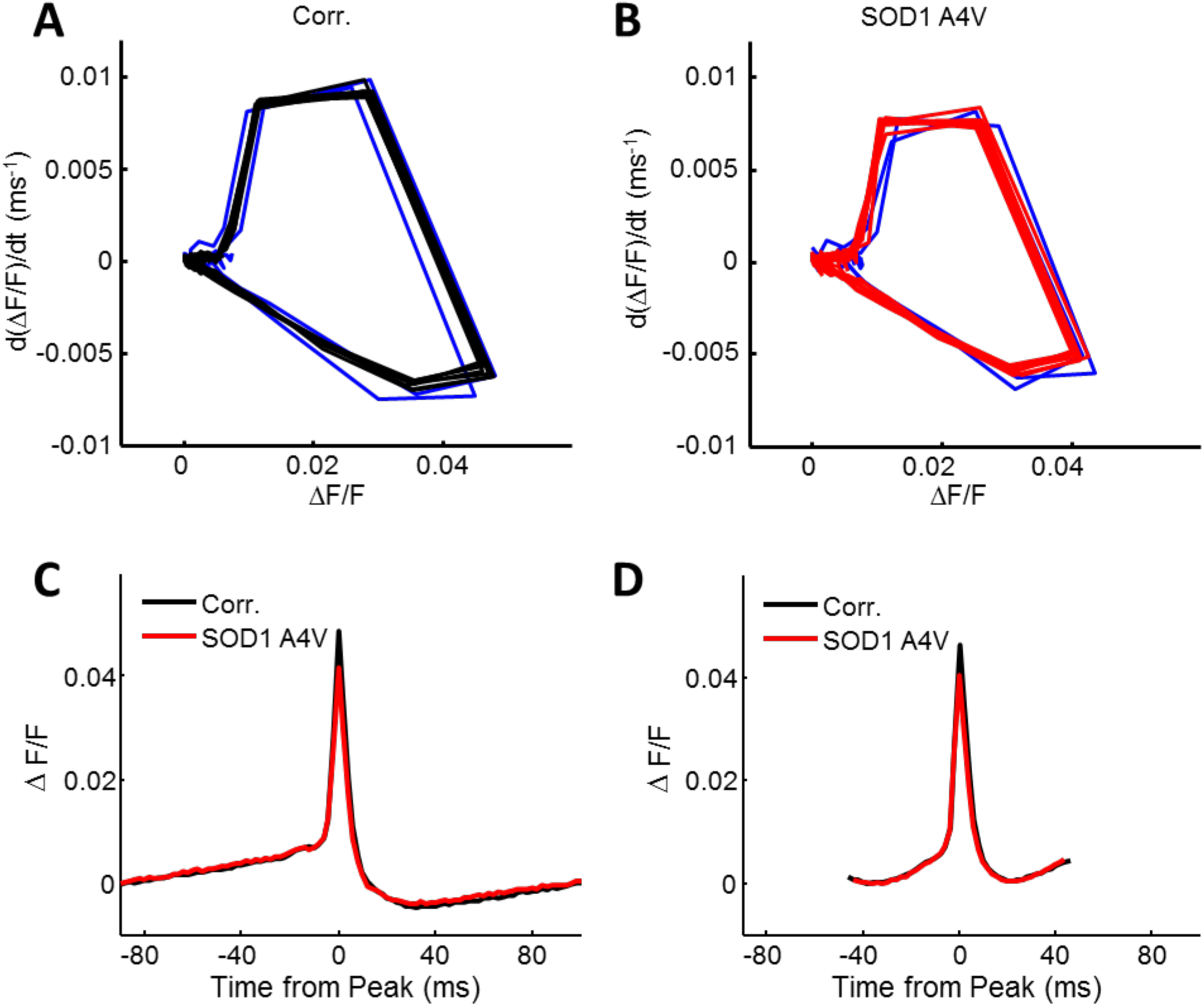
Comparison of AP waveforms in mutant and control. **(A, B)** Average phase plots, (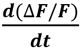 vs. Δ𝑭/𝑭), from steps with nine action potentials, for (A) Control and (B) *SOD1 A4V* cell lines. The first two spikes are highlighted in blue, after which the cells converged to a stable limit cycle. **(C)** Average action potential waveforms from all stimuli which produced **(C)** two and **(D)** nine action potentials during a single stimulus epoch. Action potentials from stimuli during which the cell entered depolarization block were excluded. Spikes were aligned in time by their peak and in Δ𝐹/𝐹 by their pre-peak minimum. Average waveforms from stimulus epochs with different numbers of action potentials are quantified in Table S2.

Finally, we studied the waveforms of individual APs. Within each 500 ms stimulus epoch, the AP waveform of each single cell was consistent from spike to spike, after the first two spikes, provided that the cell was not in depolarization block (Fig. 5; Supplemental Discussion). AP waveforms did vary with firing rate and with stimulus intensity. After controlling for these parameters, average AP waveforms still differed between mutant and control iPSC-MNs. In the weak stimulus regime, where firing rate was proportional to stimulus strength, mutants had 10% smaller AP peak amplitude than controls (*p* = 3.1 x 10^-8^ linear model coefficient t-test, significant after Holm-Bonferroni multiple comparison correction, Fig. 5, Table S2).

In accord with previous findings (Kiskinis et al., 2014), we observed statistically significant morphological differences between the mutant and control MNs. The mutants had a smaller soma and fewer projections than the controls (Fig. S6; Supplemental Methods). This observation suggested that the changes in excitability seen in the mutant cells might be a byproduct of the mutant line’s differences in morphology. We included terms for soma area and for ratio of soma area to whole-cell area (“soma fraction”) in a logistic regression model for the probability of depolarization block as a function of firing rate. When trained on control data, the coefficients on these morphological parameters were not significant (p = 0.06 for soma area, p = 0.47 for soma fraction, separate models, logistic regression coefficient t-test). When mutant data were controlled for these parameters, the difference between mutant and control remained significant (p = 1.4×10^-4^ for soma area, p = 1.2×10^-4^ for soma fraction, logistic regression coefficient t-test). Thus morphological parameters (soma area and soma fraction) did not predict the probability of depolarization block either within or between genotypes. While this analysis cannot rule out possible contributions from other morphological parameters, we focused subsequent analysis on electrophysiological variables.

### Simulations

Faced with an array of phenotypic differences and similarities between *SOD1* A4V mutant and control neurons, we sought to relate these findings to hypotheses about underlying disease mechanisms. While complex multi-compartment models of MNs have been developed (Powers et al., 2012, Powers and Heckman, 2015), it is well established that voltage recordings alone are insufficient to constrain the parameters of such models(Brookings et al., 2014). Further, considering the large cell-to-cell variability in the observed firing patterns, a morphologically and molecularly detailed model was deemed inappropriate. Instead we sought a parsimonious model which could account for multiple population-level observations with a small number of parameters.

Previous studies have singled out potassium channels in general (Kanai et al., 2006) and K_V_7 (*KCNQ*) channels in particular (Wainger et al., 2014) as a target of investigation in both *SOD1* and *C9ORF72* models of ALS; but it is not clear how these mutations affect ion channel expression or function. We asked whether changes in K_v_7 current alone could account for some or all of the observed functional effects. We performed numerical simulations of a minimal Hodgkin Huxley-type model, containing only a Na_V_ channel, a fast K_V_ channel, a slow K_V_7 channel, and a channelrhodopsin (Powers et al., 2012, Powers and Heckman, 2015)(Powers et al., 2012, Powers and Heckman, 2015)(Powers et al., 2012, Powers and Heckman, 2015)(Powers et al., 2012; Powers and Heckman, 2015). We varied the model parameters systematically and studied the resulting simulated firing patterns using the same parameters as for the experimental data.

The starting ion channel parameters were taken from previous numerical simulations of a human MN (Powers et al., 2012, Powers and Heckman, 2015), and the channelrhodopsin was modeled as a conductance with a reversal potential of 0 mV and an opening time constant of 1 ms. Following our experimental illumination protocol, simulations were run with steps of increasing maximal channelrhodopsin conductance. To account for the capacitive load from passive membrane structures and to match simulated firing rates to the range observed experimentally, we increased the membrane capacitance beyond the literature value (Supplemental Methods). The simulated cells showed stimulus-dependent firing and depolarization block, clearly recapitulating the main qualitative features of the data (Fig. 6A). The experimentally recorded waveforms varied considerably from cell to cell, so we focused on studying the dependence of spiking properties on channel conductances, rather than on trying to match simulated and experimental waveforms precisely.

**Figure 6.**
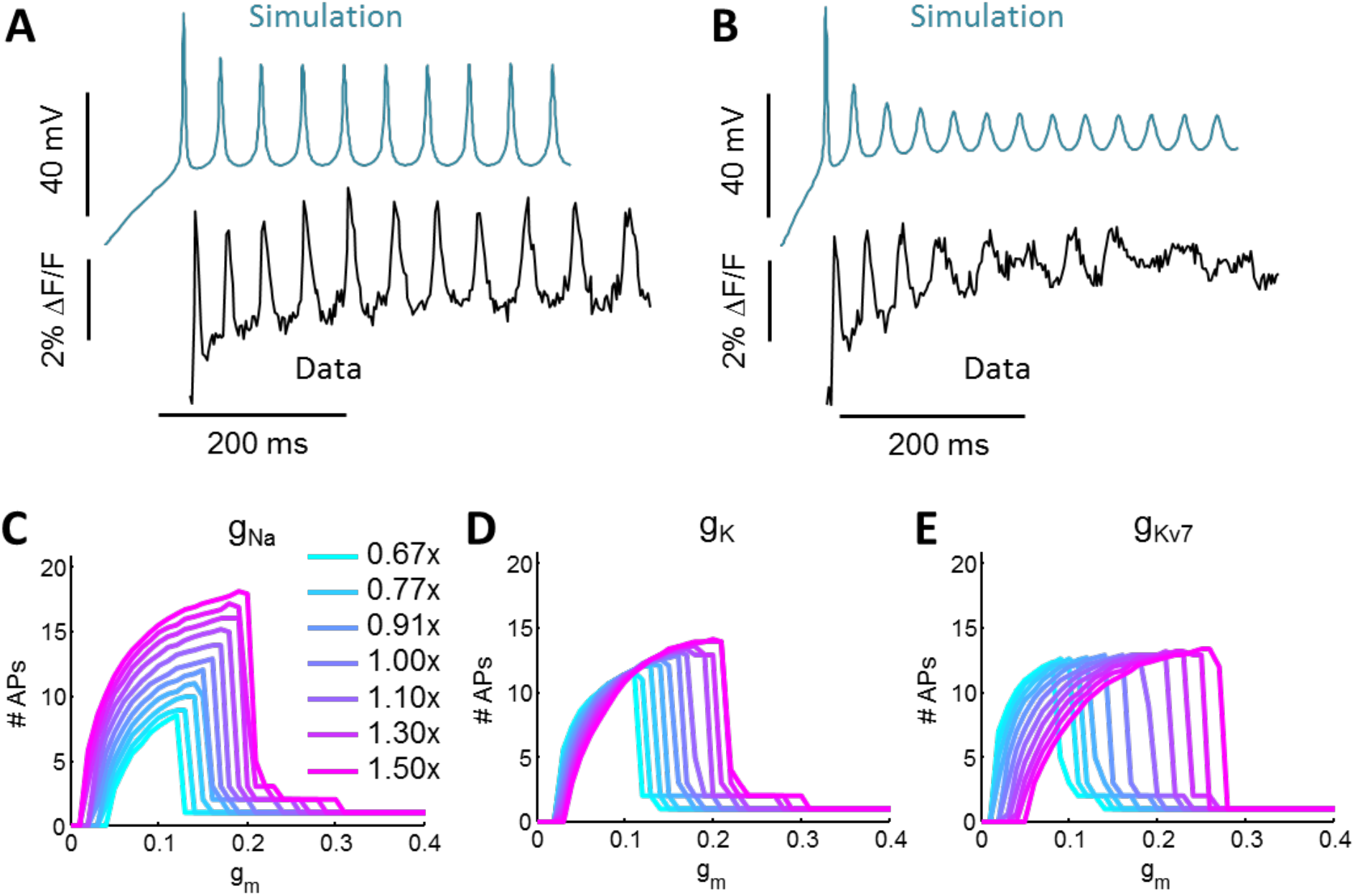
Numerical simulation of firing patterns with variable channel levels. **(A)** Simulation (blue) and fluorescence trace (black) of a neuron showing tonic firing. **(B)** Simulation (blue) and fluorescence trace (black) of the same neuron shown in (A) approaching the transition to depolarization block under strong stimulus. **(C – E)** Each trace shows the number of APs in a 500 ms interval as a function of the optogenetic stimulus strength (*g*_m_). Maximum conductances of the indicated channels were varied from to 1.5 times basal level. **(C)** Variation in Na_v_ level. **(D)** Variation in K_V_ (delayed rectifier) level. **(E)** Variation in K_V_7 level. The changes in the F-I curve that came with lower K_V_7 conductance phenotypically matched the differences and similarities between *SOD1* A4V and control neurons.

We varied each conductance in the model to determine its effect on low-stimulus excitability, threshold for depolarization block, maximum firing rate, and action potential waveform. Fig. 6 shows simulated firing rate as a function of optogenetic stimulus strength, for a range of channel conductances bracketing the original model parameters. Decreases in K_V_7 conductance increased low-stimulus excitability, decreased threshold for depolarization block, decreased spike height, and had no effect on maximum firing rate (Table S3). Thus, remarkably, changes only in the K_V_7 conductance were sufficient to reproduce all the major functional phenotypes and null results catalogued in our Optopatch experiments. Neither variation in the Na_V_ conductance nor in the fast K_V_ channel showed the correct qualitative trends (Fig. 6).

## Discussion

Our demonstration of optical electrophysiology recordings in a delicate and complex human cellular preparation opens the prospect to record large quantities of functional data in this and other human models of neuronal disease. Key to extracting meaning from these data is a statistically robust analysis pipeline and comparisons to numerical simulations at an appropriate level of detail.

### Patch vs. Optopatch

The optical and manual patch clamp techniques offer different tradeoffs in resolution and throughput, and thus should be seen as complementary rather than competing techniques. One must exercise caution in applying concepts from conventional electrophysiology to Optopatch. Principally, Optopatch and patch deliver different kinds of stimulation. CheRiff is a conductance, while electrical stimulation is via a current source. CheRiff current reverses direction when the membrane voltage crosses 0 mV, while current clamp maintains constant current irrespective of membrane voltage. In our MN simulation, the dependence of firing rate on stimulus strength and channel levels had the same qualitative features for CheRiff stimulation as for current injection. In other parameter regimes, however, the two modes of stimulation induced strikingly different firing patterns. Spike trains induced optogenetically may be a more faithful indicator of *in vivo* firing because AMPA receptors are a conductance with reversal potential of ∼0 mV, not a current source. One can simulate membrane conductances with dynamic patch clamp (Prinz et al., 2004) but this technique is not widely used.

Optical and electrical recordings also are subject to different types of noise and artifacts (Cohen and Venkatachalam, 2014). Patch clamp recordings remain the gold standard for accuracy and time resolution—one can record sub-millivolt changes in membrane voltage on a sub-millisecond timescale. One can also use voltage-clamp to dissect the contributions of distinct conductances to membrane currents. However, patch clamp measurements are slow and laborious (four cells per hour in the present experiments), increasing risk of statistical artifacts from small sample sizes in highly heterogeneous stem cell-derived cultures. Furthermore, in the most commonly used whole-cell configuration, manual patch clamp risks dialyzing cytoplasmic contents. Manual patch clamp measurements also lack spatial resolution and are exceedingly difficult to apply to a single cell on successive days.

Optopatch has lower temporal resolution (1 – 2 ms) and lower precision than patch clamp. The signal-to-noise ratio (SNR) in the optical recordings was 13.0 ± 6.7 (mean ± s.d.), corresponding to a noise level of approximately 5.2 mV in a 500 Hz bandwidth. The lower SNR relative to published results in primary rodent neurons (Hochbaum et al., 2014) was due to a smaller soma size and lower Optopatch expression in the iPSC-derived neurons. Cell-to-cell variations in QuasAr2 expression level prevented assignment of absolute voltage values to fluorescence measurements. Thus Optopatch measurements are most useful for determining spiking statistics and for examining action potential waveforms, and at present less so for quantifying sub-threshold events or absolute voltage values.

Optopatch has considerably higher throughput than manual patch clamp (34 cells per hour in the present experiments). Wide-field imaging systems could potentially increase this throughput considerably (Hochbaum et al., 2014). Although not used here, Optopatch measurements can, in principle, be readily combined with genetic or immunohistochemical targeting with cell-type-specific markers. Measurements targeted to MNs (e.g. via HB9-Cre (Peviani et al., 2012) or post-measurement HB9 staining) are a natural extension of the work. Optopatch can also probe spatial relations of electrical activity, both within and between cells.

Other neuronal measurement technologies in principle might also offer favorable tradeoffs of information content and throughput. Multi-electrode arrays (Odawara et al., 2014, Tsai et al., 2015) and nanowire devices (Robinson et al., 2012) can record non-perturbatively from neurons for long times, but cannot be genetically targeted to specific neuronal sub-types. Voltage-sensitive dyes can be fast and sensitive (Huang et al., 2015, Miller et al., 2012), but also cannot be targeted to genetically specified cell types. Ca^2+^ imaging (Pasca et al., 2011, Schöndorf et al., 2014) provides a useful surrogate measure of neuronal activity, but works best at low spike rates, does not report action potential waveforms, and is difficult to combine with optogenetic stimulation due to spectral crosstalk (Venkatachalam and Cohen, 2014). While these measurement technologies might provide similar information to Optopatch when paired with optogenetic stimulation, the challenges that will arise in these applications remain unknown until tested.

### Neuronal excitability in ALS

Our simple computational models showed non-monotonic dependence of firing rate on stimulus strength in all cases, consistent with our data. Variations in K_V_7 currents led to firing rate curves that crossed each other. Together, these observations show that “excitability” is not a well-defined attribute of a neuron, but rather depends on the magnitude of the stimulus strength. In our data, in both the mutant and controls, cells with a higher spontaneous rate were more likely to show a decrease in firing under strong stimulus: the odds of entering depolarization block increased by 34% with every 1 Hz increase in spontaneous rate (*p* = 0.005 logistic regression coefficient t-test) in control and 49% per Hz in mutant (*p* = 0.0004). Thus neurons that appeared hyperexcitable under weak or zero stimulus, tended to appear hypoexcitable under strong stimulus. One should therefore use caution in speaking of hyper-or hypo-excitability as intrinsic neuronal properties.

These observations also highlight the importance of analyzing neuronal recordings at the single-cell level, rather than simply looking at aggregate population-level statistics. The population-average curves of firing frequency vs. stimulus strength may be strikingly different from the curves for every individual neuron. Due to the nonlinear dependence of firing frequency on ion channel levels, efforts to fit the population-average data may lead to incorrect mechanistic conclusions.

The pathways by which mutations in ALS-causing genes lead to a change in neuronal excitability remain unknown. The K_V_7 potassium channel has recently emerged as a potential therapeutic target, and is the subject of an ongoing clinical trial of retigabine for ALS (McNeish et al., 2015). While expression profiling did not identify effects of the SOD1(A4V) mutation at the transcriptional level(Kiskinis et al., 2014), there are many post-translational mechanisms for regulating the K_V_7 current. This channel has multiple interaction partners (Delmas and Brown, 2005), is regulated by PtdIns(4,5)P_2_ (Suh and Hille, 2008), and is redox-sensitive (Gamper et al., 2006). Thus defects in any of these interaction partners, in lipid metabolism, or in redox homeostasis could contribute to excitability defects. Recently, deficits in trafficking of RNA granules have been found in a *TDP43* model of ALS (Alami et al., 2014). These findings provide a plausible mechanism for proteostatic deficits in ALS, including deficits in ion channels.

In a mutant *C9ORF72* model of ALS, Sareen and colleagues reported hypoexcitability of the mutants relative to controls (Sareen et al., 2013). However, the previously published plots of firing frequency as a function of stimulus strength (Fig. 3H and S12 of Sareen *et al*.) resemble our data showing depolarization block at strong stimulus (Fig. 3B, C). Though we did not probe the *C9ORF72* model, these observations raise the intriguing possibility that loss of K_V_7 conductance could provide a common mechanism across genotypes. The sufficiency of a K_V_7 deficit to account for both hyper- and hypo-excitability phenotypes cannot rule out other possible contributions to the electrophysiology, including other ion channels, shifts in channel kinetics or gating thresholds, and shifts in cellular morphology, resting potential, or ion channel spatial distribution.

The iPSC-MNs studied here represent an immature developmental state, while ALS typically strikes in adulthood. Whereas the disease-causing mutation is present throughout the life of the patient, its effect is clearly age dependent. It is not known whether time *in vitro* is a realistic proxy for chronological age *in vivo*, and if so, the relative scaling of these time-lines. Thus, while time-course data *in vitro* may yield interesting changes in function or physiology, it is not clear whether such data provide information relevant to age-related processes. Despite these limitations, functional optogenetic screening has potential uses in identifying disease mechanisms, testing prospective therapeutics, and stratifying patients.

## Acknowledgments

Scott Linderman provided an optimization algorithm for fitting piecewise linear functions. Vicente Parot helped with PCA-ICA segmentation. Samouil Farhi performed patch clamp electrophysiology measurements. This work was supported by US National Institutes of Health grants 1-R01-EB012498 and New Innovator grant 1-DP2-OD007428, and support from the Howard Hughes Medical Institute. The Kiskinis lab is supported by grants from the Les Turner ALS Foundation, Target ALS and the Muscular Dystrophy Association. EK is a Les Turner ALS Research and Patient Center Investigator. KE also gratefully acknowledges support from Project ALS, Target ALS, and National Institute of Neurological Disorders and Stroke of the National Institutes of Health under award numbers R01NS089742 and RC2NS069395.

## Materials and Methods

### Cell culture

All cell cultures were maintained at 37 °C, 5% CO_2_. Cells tested negative for mycoplasma contamination. Pluripotent stem cells were grown on Matrigel (BD Biosciences) with mTeSR1 media (Stem Cell Technologies). Culture Medium was changed every 24 hours and cells were passaged by dispase (Gibco) or accutase (Innovative Cell Technologies) as required.

### Motor neuron differentiation

Stem cell cultures were differentiated into motor neurons as previously described (Kiskinis et al., 2014). Briefly, iPSCs were dissociated to single cells and plated in suspension in low-adherence flasks (400k/mL), in mTeSR media with 10 μµ ROCK inhibitor. Media was gradually diluted (50% on day 3 and 100% on day 4) to KOSR (DMEM/F12, 10% KOSR) between days 1-4 and to a neural induction medium (NIM: DMEM/F12 with L-glutamine, NEAA, Heparin (2 μg/mL), N2 supplement (Invitrogen) for days 5-24. From days 1-6 cells were cultured in the presence of SB431542 (10 μM, Sigma Aldrich) and Dorsmorphin (1 μM, Stemgent), and from days 5-24 with BDNF (10 ng/mL, R&D), ascorbic acid (AA, 0.4 μg/mL, Sigma), Retinoic Acid (RA, 1 μM, Sigma) and Smoothened Agonist 1.3 (SAG 1.3, 1 μM, Calbiochem). On day 24 floating cell aggregates were dissociated to single cells with Papain/DNase (Worthington Bio) and plated onto poly-D-lysine/laminin-coated dishes for electrophysiological analysis. Once dissociated, MN cultures were fed every 2-3 days with complete neurobasal media (cNBM: Neurobasal with L-glutamine, NEAA, glutamax, N2 and B27), with BDNF/CNTF/GDNF (10 ng/mL, R&D) and ascorbic acid (0.2 μg/mL, Sigma).

### Gene-editing

Correction of the SOD1 A4V mutation in the ALS patient iPSC line RB9d was performed using Zinc Finger Nuclease (ZFN)-mediated targeting as described previously.(Kiskinis et al., 2014)

### Immunocytochemistry

Cell cultures were fixed in 4% PFA for 15 min at 4 °C, permeabilized with 0.2% Triton-X in PBS for 2 hours and blocked with 10% donkey serum in PBS-T (Triton 0.1%). Cells were then incubated in primary antibody overnight and secondary antibodies for 1 hour in 2% donkey serum in PBS-T after several washes in between. DNA was visualized by a Hoechst stain. The following antibodies were used: Islet1 (1:200, DSHB, 40.2D6), TUJ1 (1:1000, Sigma, T2200), MAP2 (1:10000, Abcam ab5392), GFP (1:500, Life Technologies, A10262). Secondary antibodies used (488, 555, 594, and 647) were AlexaFluor (1:1000, Life Technologies) and DyLight (1:500, Jackson ImmunoResearch Laboratories).

### Virus production

Lentivirus was produced in HEK293-T cells from previously described plasmids(Hochbaum et al., 2014) DRH334 (CamKIIα-QuasAr2, Addgene plasmid 51692) and DRH313 (CamKIIα-CheRiff, Addgene plasmid 51693). HEK293-T cells were grown in 15 cm dishes to ∼50% confluence at 37 °C and 5% CO_2_. Transfection of each gene, as well as packaging and coat proteins (psPAX2 and VSVg, respectively), was performed with poly-ethylenimine (PEI). For each 15 cm dish, a cocktail of 22 μg psPAX2, 16 μg gene, and 10 μg VSVg was suspended in 500 μL of Optimem. PEI was added to the Optimem mixture (140 μL from a 1 mg/mL stock) and vortexed briefly to mix. The mixture was incubated at room temperature for 15 minutes. After incubation, 25 mL of pre-warmed DMEM was added to the DNA/PEI mixture and gently mixed. Medium was aspirated off the 15 cm plate and the DNA/PEI/DMEM mixture was added gently onto the cells and the plate was returned to the incubator. After 48 hours, the supernatant was collected and spun for 5 minutes at 1200 g to pellet any collected cells. The supernatant was then filtered through a 0.45 μm filter, aliquoted into 1.5 mL volumes, and frozen at −80 °C.

### Gene delivery

Approximately 40,000 cells of differentiated motor neuron cultures were plated on poly-D-lysine/laminin-coated 35 mm glass bottom dishes (MatTek) for Optopatch recordings. Each dish was transduced with lentiviruses 7-10 days before scheduled recording times. A mixture of 200 μL of QuasAr2 and 70 μL of CheRiff lentiviruses were combined with 200 μL of the complete neurobasal medium, added onto the cells and left overnight in the incubator. The next morning the virus mixture was removed, plates were washed and replenished with fresh, complete neurobasal medium.

### Electrophysiology

Measurements were conducted in Tyrode’s solution containing 125 mM NaCl, 2.5 mM KCl, 3 mM CaCl_2_, 1 mM MgCl_2_, 10 mM HEPES, 30 mM glucose (pH 7.3) and adjusted to 305–310 mOsm with sucrose. Prior to imaging, neurons were incubated with 5 μM all-*trans* retinal for 30 minutes and then washed with Tyrode’s solution.

Synaptic blockers were added to the imaging medium for measurements of single-cell electrophysiology. The blockers comprised NBQX (10 μM, Tocris), D(−)-2-amino-5-phosphonovaleric acid (AP5; 25 μM, Tocris), and gabazine (SR-95531, 20 μM, Tocris). Patch clamp data was used if and only if access resistance was < 25 MΩ, and did not vary over the experiment. Recordings were terminated if membrane resistance changed by > 10%. Experiments were performed at 23 °C under ambient atmosphere.

### Optopatch recordings

Cells were imaged on a custom-built epifluorescence inverted microscope. Imaging experiments were conducted in Tyrode’s buffer (pH 7.3). Excitation of QuasAr2 was via a 500 mW 640 nm diode laser (Dragon Lasers), which provided a field of view of 31 by 37 μm, with an intensity at the sample of 800 W/cm^2^. Blue illumination from a 50 mW 488 nm solid state laser (Coherent OBIS) was modulated by an acousto-optical tunable filter (AOTF; Gooch and Housego) to control timing and amplitude of the optogenetic stimulation. The blue and red beams were then combined and imaged onto the sample through the objective lens. Images were collected with a 60x water objective (Olympus, NA 1.2) and imaged onto a scientific CMOS camera (Flash 4.0, Hamamatsu). Data was collected at a frame rate of 500 Hz. Custom code written in LabView (National Instruments) controlled the hardware.

The stimulation protocol consisted of:

1. 10 s of continuous red light to measure spontaneous firing
2. 50 ms with red light only
3. 500 ms with red and blue light to stimulate firing
4. 500 ms with red light only
5. 5 s with no light for cell recovery

Steps 2-5 were repeated 8-10 times with increasing intensities of blue light.

Data Analysis and Simulation procedures are described in detail in the Supplemental Experimental Methods.

## Supplemental Material

**Figure S1.**
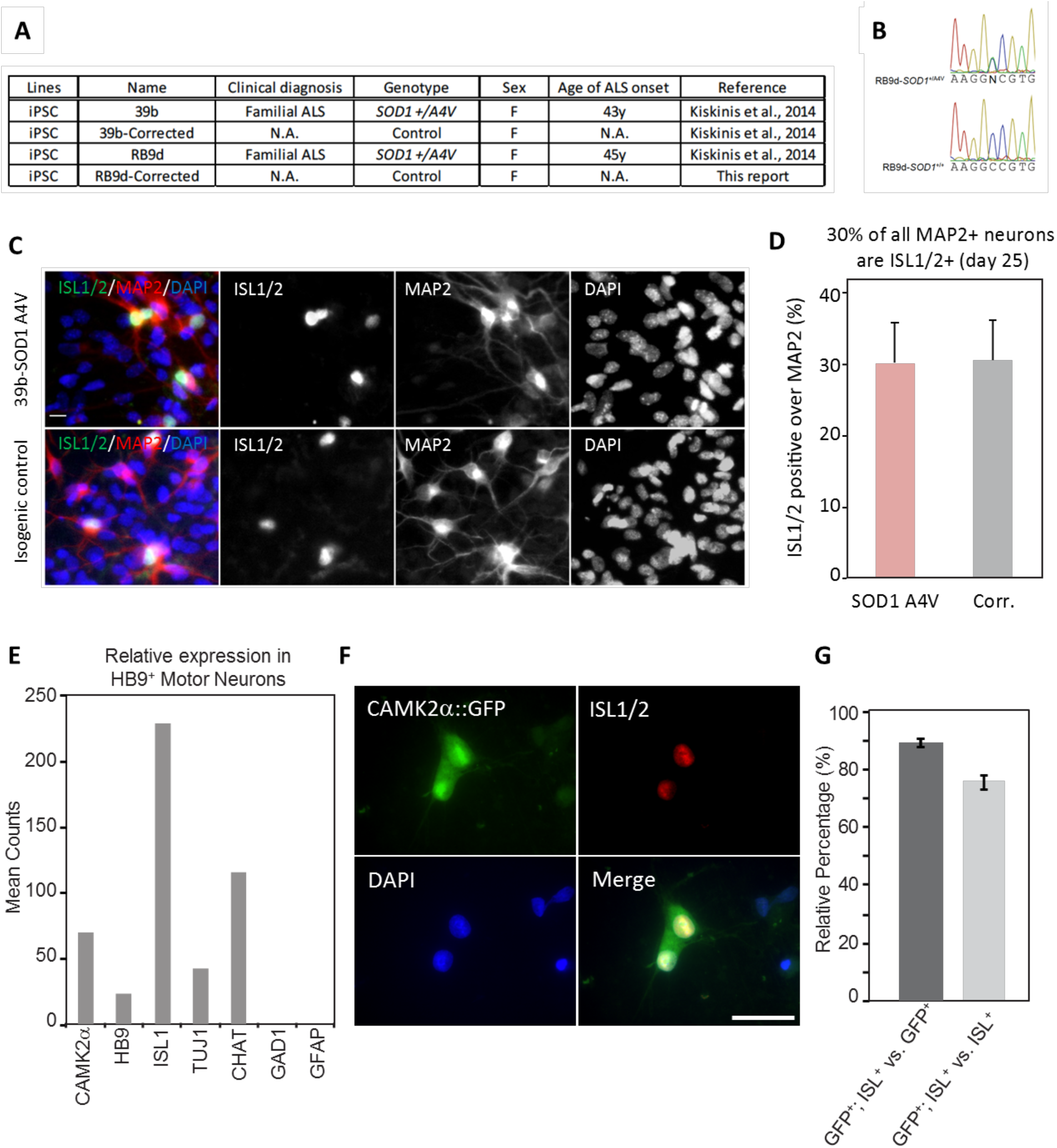
Characterization of human motor neurons derived from iPSCs. Related to Fig. 1. (A) ALS patient-iPSC lines and isogenic controls used in this study. (B) Sequencing chromatographs of exon 1 of SOD1 in patient line RB9d before and after gene-targeting demonstrate the successful correction of the A4V mutation. Editing was performed using Zinc Finger Nucleases as described previously (Kiskinis et al., 2014). (C) Immunocytochemistry for a MN marker (ISL1/2), a pan-neuronal marker (MAP2) and a nuclear marker (DAPI). Scale bar 50 μm. (D) Quantification of MN populations. Error bars represent standard deviation of the percentage of ISL1/2^+^ cells from 4 images from 1 differentiation (SOD1 A4V: *n* = 407 MAP2^+^ cells; Corrected: *n* = 291 MAP2^+^ cells). (E) Analysis of previously published RNA-Seq data from iPSC-derived HB9^+^ MNs purified via FACS (Kiskinis et al., 2014; GEO entity: GSE54409). These cells expressed CAMK2α, HB9, TUJ1 and CHAT, while the glial marker GFAP and the interneuron marker GAD1 were absent. (F) Immunocytochemistry to detect the selectivity and specificity in MNs of lentivirally delivered constructs with the CamKIIα promoter. Images show MN cultures infected with lentivirus containing CamKIIα-driven eGFP. Cells were labelled by immunocytochemistry with antibodies for ISL1/2 and eGFP, and DAPI; scale bar 30 μM. (G) Quantification of images as in (E). 89% of eGFP^+^ cells were also ISL^+^. 75% of all ISL^+^ MNs were also eGFP^+^. Error bars represent s.e.m. from 42 images taken from 2 independent differentiations, *n* = 1147 ISL^+^ hiPSC-MNs.

**Figure S2.**
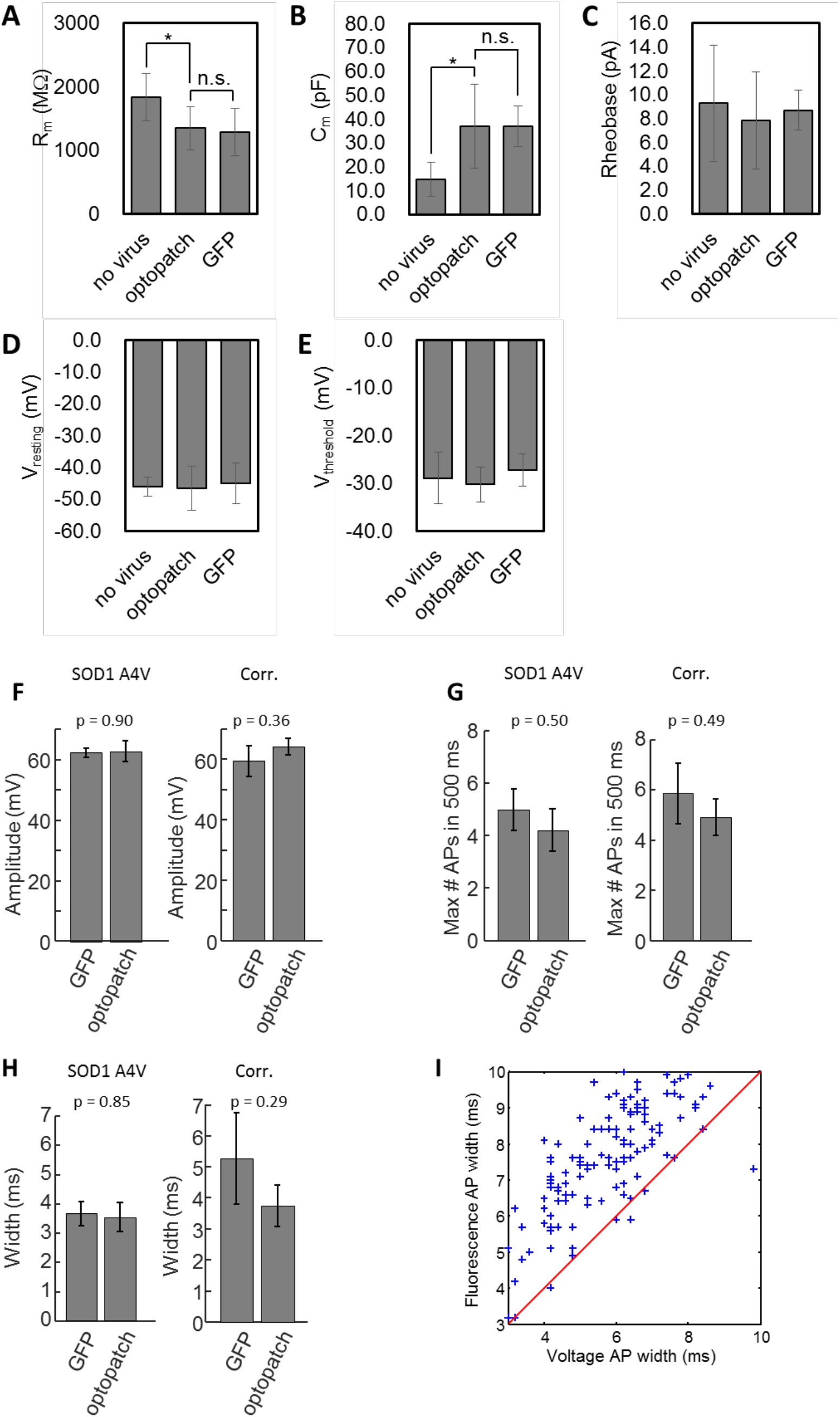
Validation of Optopatch measurements in iPSC-MNs. Related to Fig. 2. **(A-E)** Effect of Optopatch expression on baseline electrophysiology. Comparisons between iPSC-MNs expressing no lentivirus, Optopatch constructs, or an eGFP control showed no significant differences between Optopatch and eGFP expression in **(A)** membrane resistance, **(B)** membrane capacitance, **(C)** rheobase current, **(D)** resting potential, and **(E)** action potential initiation threshold (See Table S1). The CaMKIIα promoter targeted expression (of either eGFP or Optopatch) to more mature, and hence larger MNs (Prè et al., 2014), leading to smaller membrane resistance and larger membrane capacitance compared to non-transduced cells. Error bars represent s.e.m.. Measurements were on *n* = 10 – 14 cells per condition. **(F – H)** Comparison of spiking parameters in iPSC-MNs expressing either Optopatch constructs or an eGFP positive control, as recorded by manual patch clamp. Current was injected in 500 ms pulses of nineteen amplitudes (5 pA to 95 pA; n = 8 39b-C cells with eGFP, n = 15 39b-C cells without Optopatch, n = 9 39b cells with eGFP and n = 9 39b cells with Optopatch). **(F)** Within each genotype, cells that expressed eGFP and those that expressed Optopatch had indistinguishable average action potential amplitude (p = 0.90 for 39b, p = 0.36 for 39b-C, unpaired t-test). **(G)** The cells reached indistinguishable maximum firing rates (p = 0.50 for 39b, p = 0.49 for 39b-C, unpaired t-test). **(H)** The width of the spikes at rheobase was indistinguishable (p = 0.85 for 39b, p = 0.29 for 39b-C, unpaired t-test). **(I)** Comparison of action potential width measured optically vs. electrically. Optical measurements reported an AP width that was greater than the electrically recorded width by 1.8 ± 1.1 ms (mean ± s.d.; *n* = 148 spikes, 4 cells).

**Figure S3.**
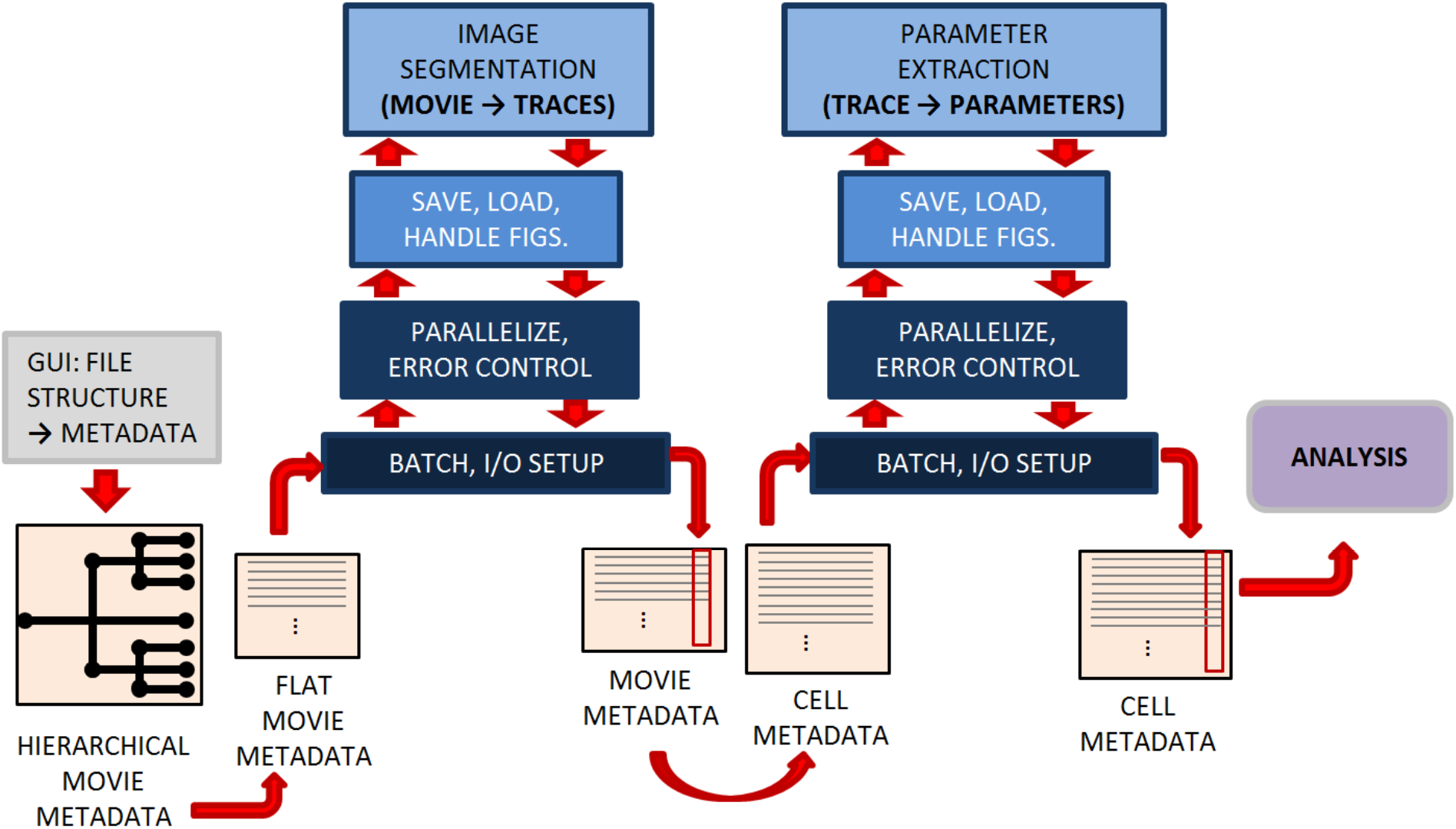
Data pipeline for image segmentation and spike parameterization. Related to Fig. 2. Metadata from each experiment and sub-experiment is created using a graphical user interface. The hierarchically organized metadata is then flattened into a matrix listing each recording and its associated metadata. This organization facilitates parallel analysis on a computer cluster. In the first stage of analysis, movies are translated into a set of fluorescence traces via image segmentation; morphological and expression level measurements are also recorded. Summary figures of each movie’s analysis are created and data from each movie’s analysis is saved individually. The metadata is then broken up by cell (rather than by movie) and sent to the parameter extraction pipeline which again is parallelized. In this stage, each cell’s fluorescence trace is loaded and parameterized. A summary figure is created and again the results are saved individually. The final list of cells and their parameterizations can then be loaded for further analysis. The entire pipeline is implemented in MATLAB and run on the Odyssey Research Computing cluster at Harvard University.

**Figure S4.**
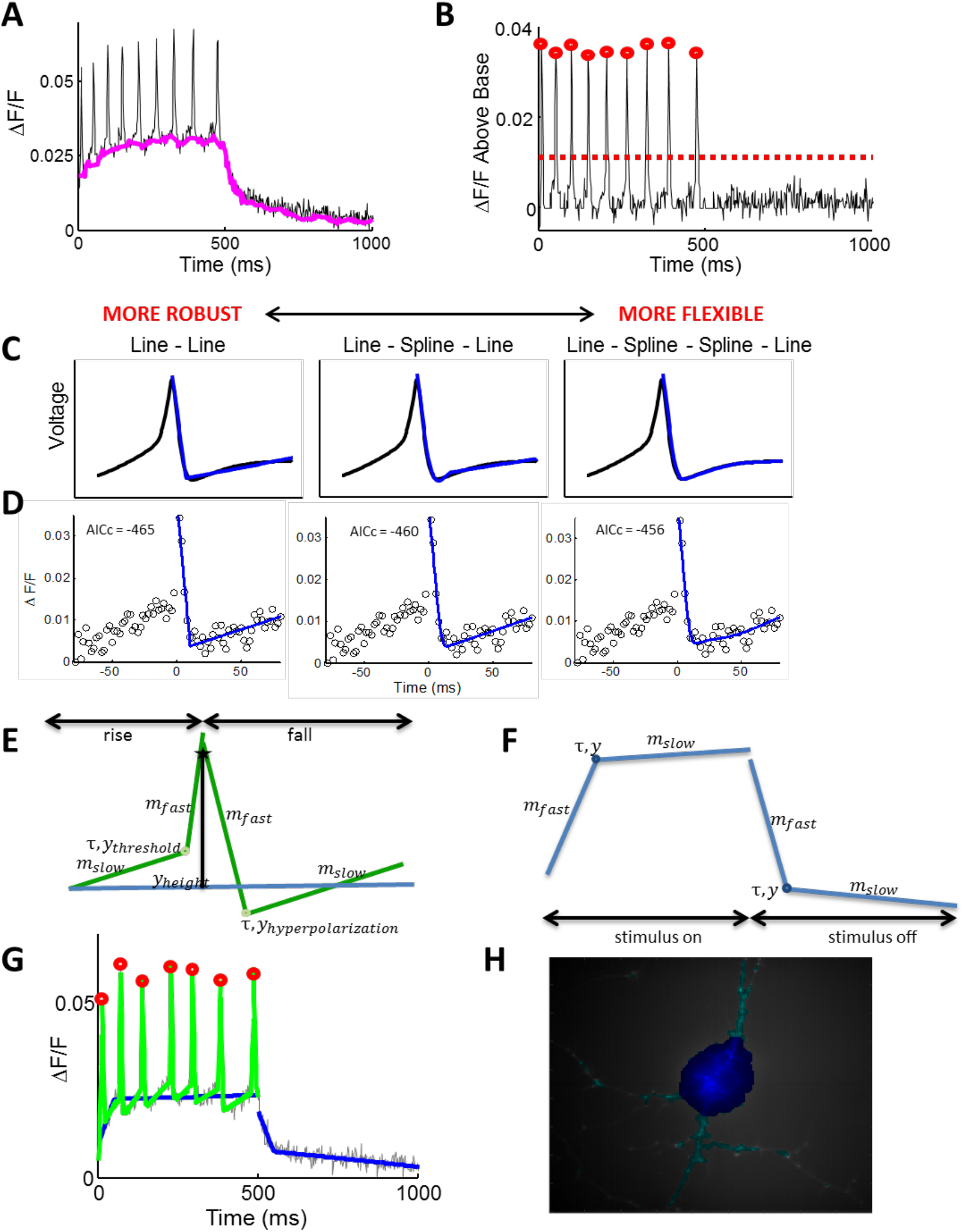
Data reduction and parameterization of spiking waveforms. Related to Fig. 2. **(A)** Fluorescence traces were temporally filtered with a specialty ordinal filter to separate spikes from baseline (window size of 20 ms, 40^th^ percentile taken as baseline estimate). **(B)** A threshold for spike identification was set based on an estimate of the noise. **(C – F)** Information-theoretic determination of optimal parameterization of optically recorded spike waveforms. We created a hierarchy of models with different number of parameters; adding more parameters increased flexibility but decreased robustness to noise. The least complex model of the AP downstroke consisted of two lines meeting at a point. The next model added one spline joining the two lines. The most flexible model contained two splines. **(C)** Fit of each model to an AP waveform recorded by manual patch clamp.(Bean, 2007) **(D)** Fit of each model to single-trial Optopatch data. Corrected Akaike’s information criterion (AICc) balances the quality-of-fit with the complexity of the model. Lower scores indicate a better model for the given data. In this example (as in 95% of tests), AICc was lowest for the least complex and most robust two-line model. **(E)** Parameterization of optically recorded action potential waveforms **(F)** and baseline waveforms. **(G)** Final parameterization, consisting of the fitted baseline and action potential models. **(H)** Automated identification of soma (dark blue) and processes (cyan) calculated from the activity-based map of the single-cell profile.

**Figure S5.**
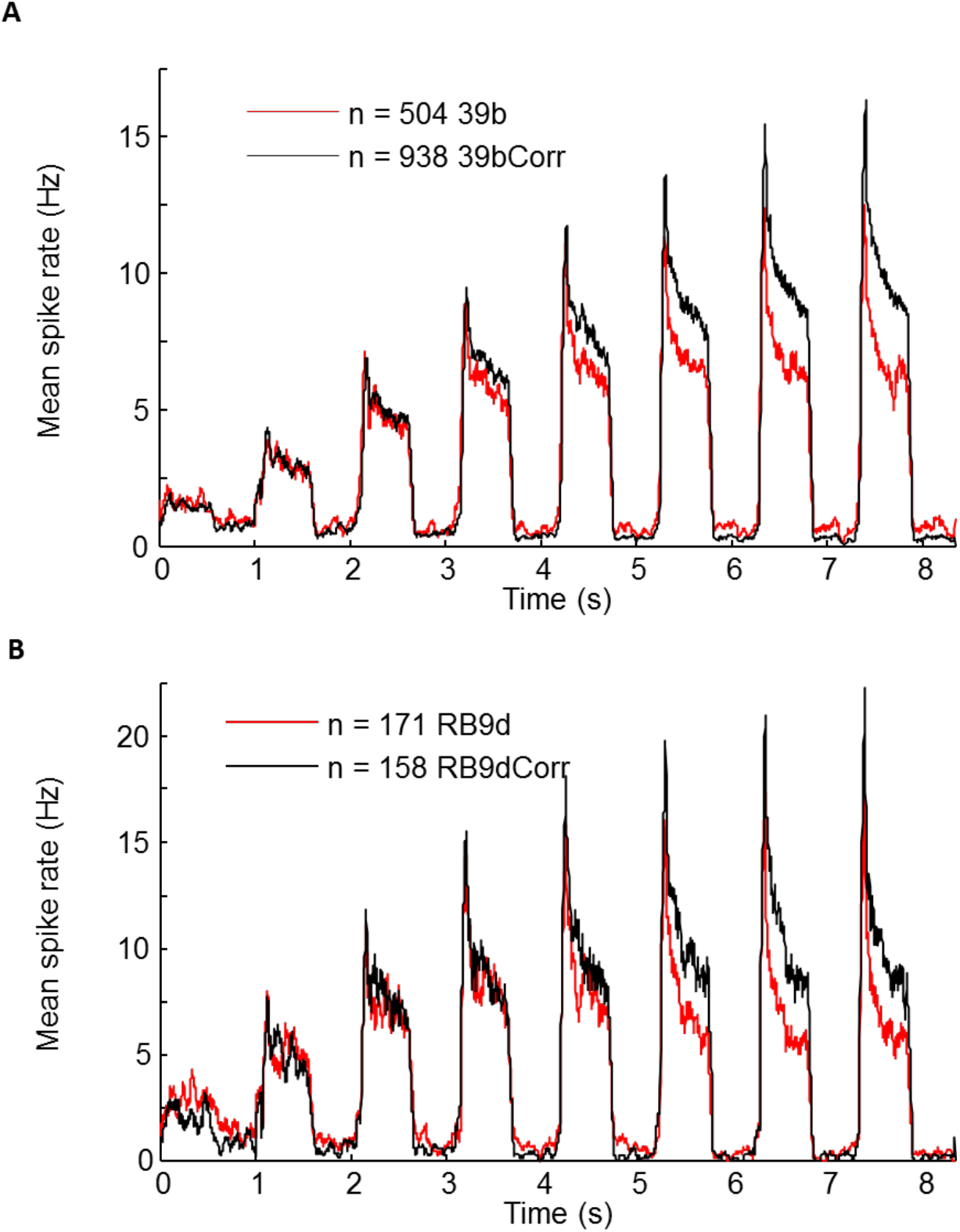
Replication of core experimental results in two distinct isogenic pairs. Related to Fig. 3. A) The 39b SOD1(A4V) line and its isogenic corrected pair. B) The RB9d SOD1(A4V) line and its isogenic corrected pair. As in the 39b cell line, the SOD1 (A4V) cells showed hyperexcitability under weak stimulus, but hypo-excitability under strong stimulus.

**Figure S6.**
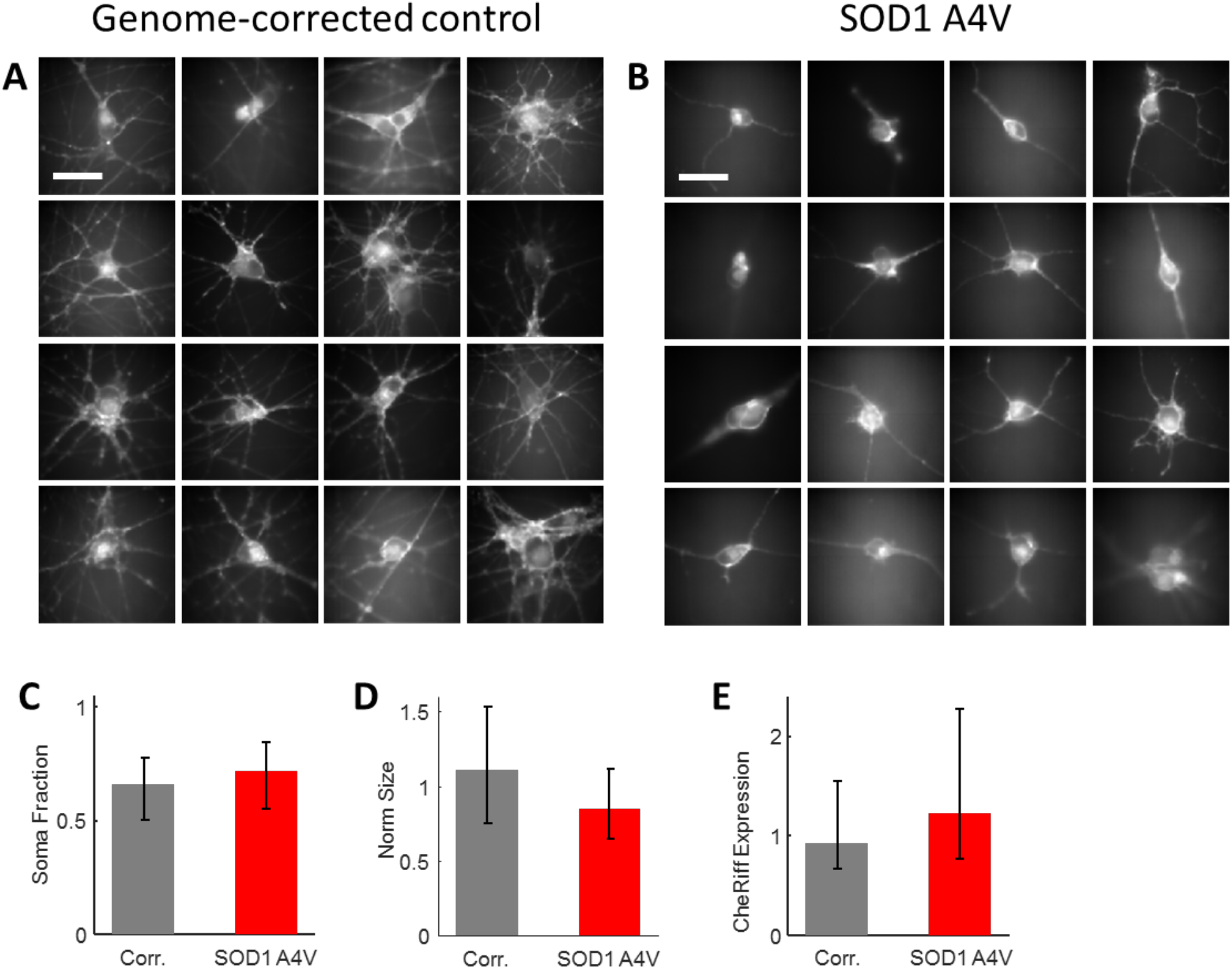
Morphological differences between control and SOD1 A4V motor neurons. Related to Fig. 3. Images show **(A)** corrected control and **(B)** *SOD1* A4V mutant iPSC-MN. Fluorescence is from QuasAr2. Scale bars in (A) and (B) 20 μm. **(C)** Median fraction of the cell area occupied by soma (as opposed to dendrite) in the control and mutant iPSC-MNs; error bars show the 25^th^ and 75^th^ percentiles of the population. The difference between the two populations was significant to p = 1×10^-10^ (0.66 in control, 0.72 in mutant, unpaired t-test). **(D)** Median size of the control and mutant cell somas, normalized to the population median; error bars show 25^th^ and 75^th^ percentile. The difference between the two populations was significant to p = 8×10^-17^ (1.11 a.u. in control, 0.84 in mutant, unpaired t-test). **(E)** Difference in median CheRiff expression level between control and mutant cells, normalized to the whole population median. Error bars shows the 25^th^ and 75^th^ percentile. The difference between the two populations is significant (0.92 a.u. in control, 1.22 in mutant, p = 2×10^-5^, unpaired t-test). In C – E, control *n* = 843 cells, mutant *n* = 331 cells.

**Table S1.**
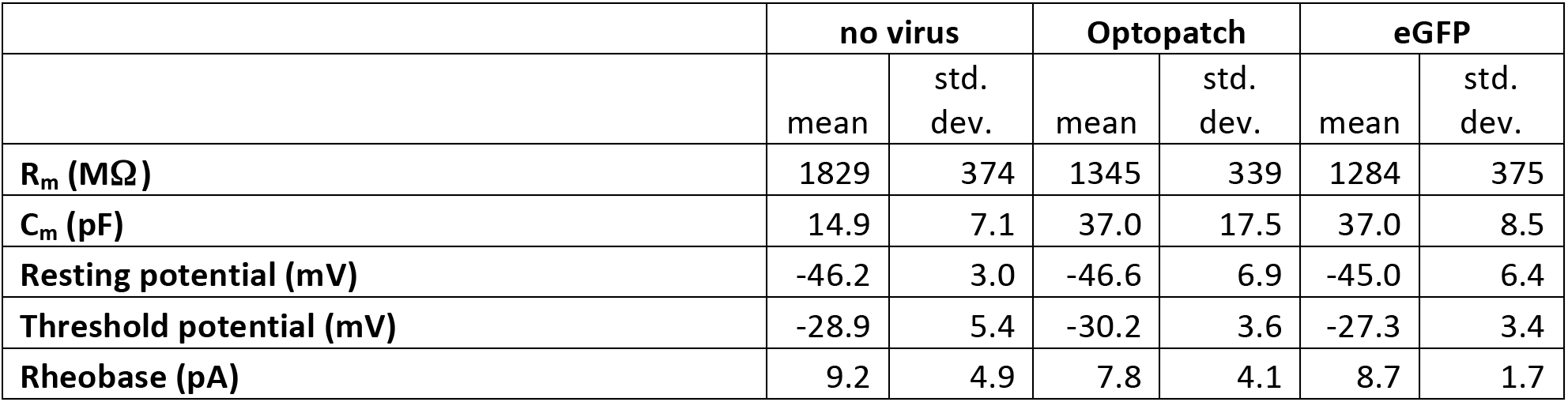
Comparison of electrophysiological parameters of iPSC-MN infected with either no virus, CaMKIIa-driven Optopatch, or CaMKIIa-driven eGFP. The data are shown graphically in Fig. S2.

|  | no virus |  | Optopatch |  | eGFP |  |
| --- | --- | --- | --- | --- | --- | --- |
|  | mean | std. dev. | mean | std. dev. | mean | std. dev. |
| **R<sub>m</sub> (MΩ)** | 1829 | 374 | 1345 | 339 | 1284 | 375 |
| **C<sub>m</sub> (pF)** | 14.9 | 7.1 | 37.0 | 17.5 | 37.0 | 8.5 |
| **Resting potential (mV)** | -46.2 | 3.0 | -46.6 | 6.9 | -45.0 | 6.4 |
| **Threshold potential (mV)** | -28.9 | 5.4 | -30.2 | 3.6 | -27.3 | 3.4 |
| **Rheobase (pA)** | 9.2 | 4.9 | 7.8 | 4.1 | 8.7 | 1.7 |

**Table S2.**
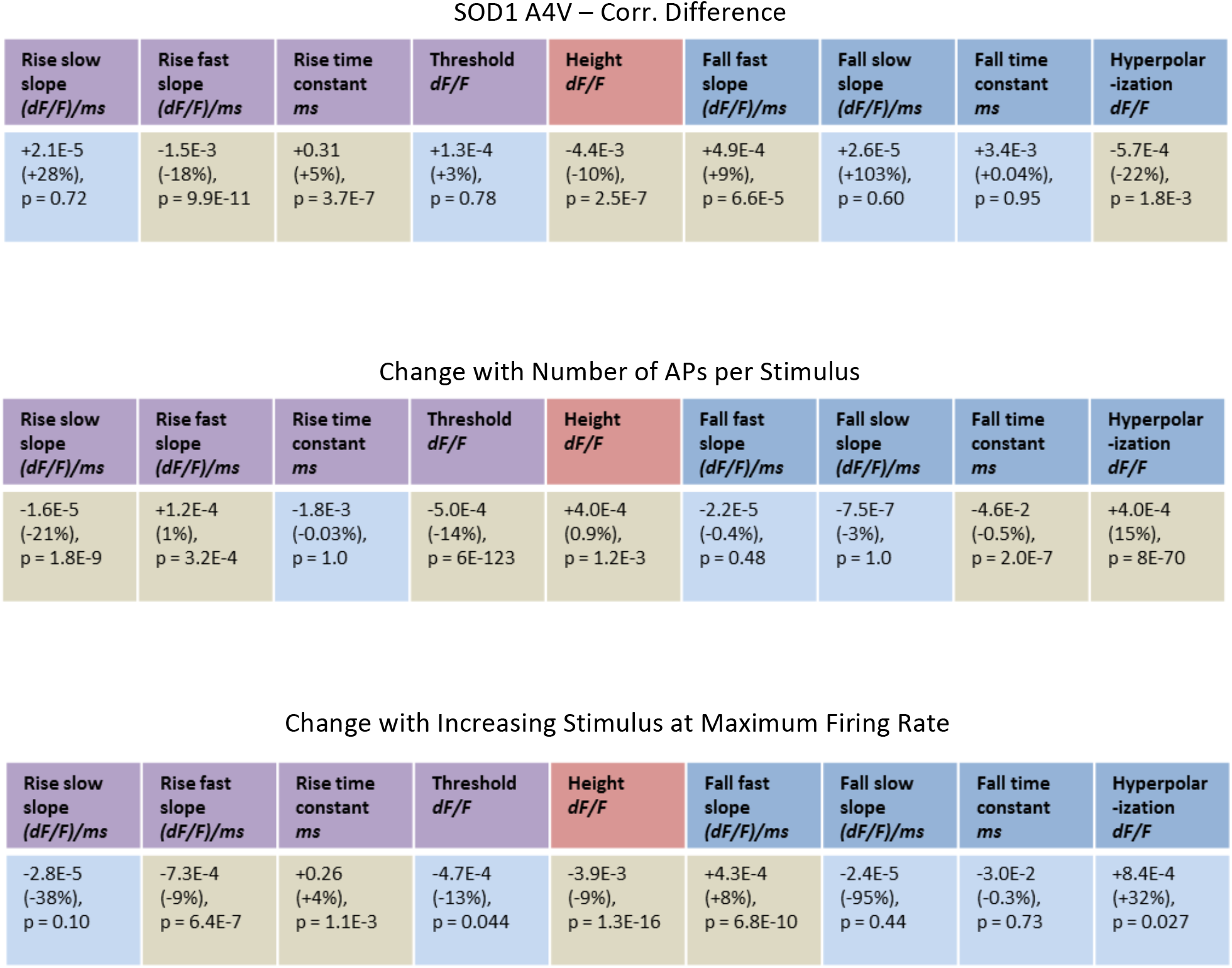
Changes in action potential waveform parameters between *SOD1* A4V and genome corrected **controls** (see Figure S5 for definitions). We performed a linear regression to determine the factors that influenced average waveform parameters for each stimulus and each cell, up to the stimulus at which the maximum firing rate first occurred. The control parameters were: cell line (mutant or control, a categorical variable), number of action potentials in the stimulus epoch, stimulus strength (to control for blue light-induced photoartifacts), and dish recording day (to control for day-to-day variability). Each column of the **SOD1 A4V - Corr. Difference** table lists the regression coefficient for genotype (mutant vs. control), the percent change relative to the control’s average, and the p value from the regression coefficient t-test. Coefficients are highlighted in olive-green which are significant to an α = 0.01 threshold after Holm-Bonferroni multiple hypothesis correction (9 hypotheses from the 9 parameters).

**Table S3.** Change in action potential waveform in numerical simulations with increases in channel conductance. For each channel or set of channels, simulations were run with a range of conductances and a range of optogenetic stimulus strengths corresponding to changes in channelrhodopsin conductance (𝒈_𝒎_, see Fig. 6). The simulated voltage trace was down-sampled to match the data acquisition frame rate (500 Hz), and action potentials were fit with the same set of parameters used in the fluorescence data. We then fit a linear regression on the average waveform parameter from each step with control variables for the channel conductance and the input conductance. Rows of the table show the regression coefficient (in units of waveform parameter per base simulation conductance) for variations in K_V_7, delayed rectifier K_V_, and Na_V_. The coefficient as a percentage of the base simulation average is shown in parenthesis. As another indicator of the strength of the relationship we also show the coefficient as a fraction of its standard deviation, 𝛽 (we can write 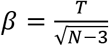 where 𝑇 is the t-statistic and 𝑁 − 3 is the number of degrees of freedom in our regression).

|  | Rise slow<br>slope<br>(dF/F)/ms | Rise fast<br>slope<br>(dF/F)/ms | Rise time<br>constant<br>ms | Threshold<br>dF/F | Height<br>dF/F | Fall fast<br>slope<br>(dF/F)/ms | Fall slow<br>slope<br>(dF/F)/ms | Fall time<br>constant<br>ms | Hyperpolar-<br>ization<br>dF/F |
| --- | --- | --- | --- | --- | --- | --- | --- | --- | --- |
| K<sub>v7</sub> | +4.4E-5<br>(+35%)<br>$\beta = 0.20$ | +1.7E-3<br>(+15%)<br>$\beta = 0.24$ | -7.0E-1<br>(-12%)<br>$\beta = -0.20$ | +2.2E-3<br>(+132%)<br>$\beta = 0.68$ | +6.0E-3<br>(+11%)<br>$\beta = 0.49$ | -1.5E-3<br>(-15%)<br>$\beta = -0.23$ | +1.0E-5<br>(+25%)<br>$\beta = 0.05$ | -3.2E-1<br>(-5%)<br>$\beta = -0.09$ | -1.5E-3<br>(-109%)<br>$\beta = -0.63$ |
| K<sub>dr</sub> | +1.6E-4<br>(+128%)<br>$\beta = 0.67$ | +1.4E-3<br>(+13%)<br>$\beta = 0.18$ | -8.4E-1<br>(-15%)<br>$\beta = -0.23$ | +4.2E-3<br>(+253%)<br>$\beta = 1.26$ | +3.6E-3<br>(+6%)<br>$\beta = 0.29$ | -5.7E-3<br>(-57%)<br>$\beta = -0.96$ | +1.9E-4<br>(+468%)<br>$\beta = 0.78$ | -2.4<br>(-39%)<br>$\beta = -0.81$ | -3.8E-3<br>(-267%)<br>$\beta = -1.35$ |
| Na | +8.7E-5<br>(+67%)<br>$\beta = 0.10$ | +2.4E-2<br>(+211%)<br>$\beta = 1.81$ | -5.8<br>(-102%)<br>$\beta = -1.13$ | +1.9E-3<br>(+117%)<br>$\beta = 0.30$ | +4.1E-2<br>(+73%)<br>$\beta = 2.44$ | -1.6E-2<br>(-165%)<br>$\beta = -1.76$ | +9.9E-6<br>(+24%)<br>$\beta = 0.01$ | -4.6<br>(-74%)<br>$\beta = -1.05$ | +8.5E-4<br>(+60%)<br>$\beta = 0.16$ |

## Supplemental Discussion

### Optical crosstalk in Optopatch measurements

The fluorescence signal showed a slow increase in baseline during each optical stimulus epoch (Fig. 1D). This increase did not occur in patch clamp recordings with optical stimulation, indicating that the increase did not correspond to a real increase in membrane potential. The increase also did not occur in optical recordings with stimulation via current injection from the patch pipette, indicating that the increase was not a slow response of the voltage indicator to membrane depolarization. The increase only occurred with simultaneous blue light stimulation and fluorescence imaging. These observations implied that the increase in baseline fluorescence was due to blue light photo-production of a fluorescent product. This effect was not observed in prior experiments with rodent primary neurons due to higher CheRiff expression in the rodent cells (and consequently lower blue light intensities), and better membrane trafficking of QuasAr2 in the rodent cells.

### Spike parameterization and model reduction

In functional fitting, choosing a good functional form is critical. On the one hand, it must be capable of fitting the data well and providing a meaningful description of relevant parameters; on the other hand, it must be robust and avoid over-fitting. We first attempted to fit action potential data with exponential rise and decay functions, but the fit was often qualitatively poor. We found more success with combinations of lines and splines. For instance, to describe the shape of an action potential after its peak, one line could be used to fit the fast action potential down-stroke, a connected spline segment to model the after-hyperpolarization, and then another connected line to fit the return to baseline (Fig. S4).

To address the problem of robustness and overfitting we employed an information-theoretic model reduction scheme. Our line-spline-line functions lent themselves well to the creation of a hierarchy of models with varying degrees of complexity (Fig. S4). We tested three models, with zero, one, or two splines between two lines. The point at which each segment (line or spline) joined with the next was allowed to vary along both the ΔF and time axes, but continuity between segments was enforced. These models had four, six and eight parameters respectively: the slope of the two lines and position of one, two, or three joining points.

To balance the tradeoff between quality-of-fit and model complexity we employed corrected Akaike information criteria (AICc).(Burnham and Anderson, 2003) This criterion enabled us to choose the model with the best level of complexity for our data, based on how closely the data matched the model, how many parameters must be fit in the model, and how many data points are being fit.

Our likelihood function assumed Gaussian noise in ΔF; the width of the distribution was estimated by subtracting a median filtered copy of each cell’s recording (to remove baseline fluctuations) and then taking the 50^th^ minus the 16^th^ percentile in the residual. We fit a sample of 200 action potentials randomly selected from 50 cells using the three models. Although these functional models describe action potential shape well both pre- and post-peak, we focused on post-peak since after-hyperpolarization is in general more difficult to describe. The results from AICc showed that the simplest model (0 splines) was optimal in 95% of sample action potentials, versus 3% for the one-spline model and 2% for the two-spline model. We therefore used the simplest model. We fit the pre- and post-peak sections of each action potential separately to a pair of lines which could vary in their slope and the point at which they met (Fig. S4). We called this model a “variable hinge.”

### AP waveform statistics

We studied action potential waveforms in the pre-depolarization block regime. Here, after the first two spikes upon stimulus onset, cells typically settled into a consistent limit cycle with little change in action potential waveform over the course of the stimulus (Fig. 5). We therefore averaged together the parameters of the waveforms of each action potential after the first 100 ms of stimulation.

As action potential firing rate increased, action potential waveforms changed. We constructed a linear model for each average action potential waveform parameter from each step stimulus, with a control coefficient for the number of action potentials in the step from which the waveform was taken. To control for any residual variation caused by blue light crosstalk, we also included a control coefficient for the stimulus number in our linear model. To study differences between control and mutant lines, we added a categorical variable to the model for cell line (this variable was non-interacting with the others). The model was fit to a dataset containing waveform parameters from every step in every cell up to and including the first step at which the maximum firing rate was reached. In reporting parameters that changed significantly between the two populations, we applied the Holm-Bonferroni method to correct for multiple hypothesis testing (see Methods). Regression coefficients and p values are presented in Table S2.

We compared the set of differences and similarities between action potential waveforms from mutant and control cells to other axes of variability between action potential waveforms. The set of genotype-dependent changes was different from those which occurred as the firing rate increased (Table S2). Genotype-dependent changes were nearly identical, however, to the set of changes which occurred as one moved from the first stimulus at which the maximum firing rate was reached to the last stimulus at which the maximum firing rate occurred before depolarization block (Table S2). This observation suggests that the differences in waveform between control and mutant cell lines may come from the same mechanism that produced differences in the probability of depolarization block.

## Supplemental Experimental Procedures

### Data analysis

#### Data cleaning

Under external synchronous triggering, the Flash 4.0 camera rounds the exposure time to the nearest 10 μs. This variation in exposure time is inconsequential for long exposures, but led to spurious noise of 0.5% at an exposure time of 2 ms, due to the asynchronicity of the computer clock triggering the camera and the camera’s internal clock. We used the whole-field image intensity to estimate this rounding error and then divided the pixel values in each frame by the estimated exposure time to correct for the variation in exposure time.

### Image processing and segmentation

Our image segmentation pipeline was adapted from that of Mukamel and coworkers (Mukamel et al., 2009) and consisted of pre-process filtering, PCA-ICA in the time domain, and post-process statistical analysis. A key technical challenge in the analysis was that each pixel had a low signal-to-noise ratio (SNR) due to the short exposure time (2 ms) and the low intrinsic brightness of QuasAr2. Noise from neighboring pixels was uncorrelated, while true signals from neighboring pixels tended to be correlated because each cell extended over many pixels. We thus started the analysis by performing median spatial filtering to improve the per-pixel signal-to-noise ratio. This procedure also removed the effect of sparse bad pixels in the camera image sensor.

A second key challenge was that the dominant fluorescence dynamics corresponded to (a) whole-field photobleaching, and (b) stepwise increases and decreases in whole-cell fluorescence triggered by the blue stimulus illumination. Both of these sources of temporal variation were highly correlated between cells and thus were not good signals for activity-based image segmentation. Single-cell spiking patterns, however, were statistically independent between cells and provided a robust segmentation signal. We therefore applied high-pass filtering in the time domain with a mean-subtracted Gaussian filter set to accentuate action potentials (window size of 20 ms, Gaussian with standard deviation of 3 ms).

A third key challenge was that the signals of interest were sparse in space and time, while the noise was broadly distributed. The PCA-ICA protocol involved calculations of cross-correlation functions. Inclusion of noise-dominated elements of the data in these calculations led to increases in noise without improvements in beneficial signal. We therefore chose only to apply PCA-ICA to the region in time corresponding to our staircase stimulation, rather than including the spontaneous imaging region, since most activity occurred during stimulation.

Using the notation 𝐴_*x,y*_ for matrices where *x* and *y* represent the domains a matrix maps *from* and *to* respectively, these last two steps translated a movie matrix 𝑀_*r,t*_ (where *r* is the pixel domain and *T* is the time domain) into a high-pass-filtered and abbreviated movie matrix 𝐻_*r,t*_ (where *t* is the abbreviated time domain).

There were fewer frames than pixels in our movies, so we employed time-domain-covariance matrix PCA to reduce dimensionality of the data. The covariance matrix 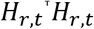 was decomposed into its eigenvectors 𝐸 with eigenvalues in the diagonal matrix 𝐷; the twenty with the largest eigenvalues were retained to form the eigenvector/principal component matrix 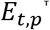 (where *p* is the principal component index). We applied ICA to the matrix 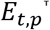 using the FastICA algorithm with a symmetric approach. The expected distribution of fluorescence values for each pixel was a Gaussian (noise) with a one-sided long tail (action potentials). This distribution has high skew and so we employed the function 𝑔 𝑢 = 𝑢^3^ as a contrast function.(Hyvärinen and Oja, 2000) Six independent components, 𝐶_*t,v*_, were calculated, but these did not correspond to true neuronal activity waveforms because they were extracted from the temporally high-pass-filtered movie.

ICA produced a separation matrix 𝑆_*s,v*_ which mapped from the principal component domain to the ICA temporal voices domain (*v*), i.e. 𝐶_*t,v*_= 𝐸_*t,p*_𝑆_*p,v*_, We generated a spatial filter 𝐹_*s,v*_ which mapped from the spatial pixel domain to the spatial voices domain:

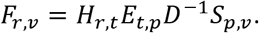

Application of this filter to each frame of the original movie provided a final fluorescence trace, T_*v,t*_, for each cell in the movie, i.e. 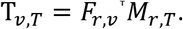

Once the traces were constructed, we performed spike finding (see below). Traces with five or fewer spikes in the entire recording were considered to be either noise or inactive cells and were discarded. A common failure mode for ICA produced traces with positive spikes on top of an inverted baseline. These were also automatically discarded by checking for negative changes in baseline during high stimulus.

We also extracted an image of each cell, 𝐼_*r,v*_, by cross-correlating the high-pass-filtered time-trace of each cell with the high-pass-filtered movie, i.e. 𝐼_*r,v*_ = 𝐻_*r,t*_𝐶_*t,v*_. Cells may be partially overlapping but they are still spatially sparse in the sense that they typically only cover a relatively small region of the whole frame. The distribution of values in 𝐼_*r,v*_ comprised Poisson-distributed background noise, with a long tail corresponding to the cell. We set a dynamic threshold of 1.8 times the estimated Poisson parameter λ to determine which pixels were most likely to be on-cell. The final cell mask for a given cell 𝑣_*j*_ was then given by 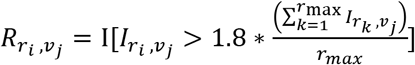 where I is the indicator function and 𝑟_*max*_ is total number of pixels.

### Morphological analysis and calibration of ΔF/F

The PCA-ICA method described above produced traces which were scaled and offset versions of the true fluorescence trace; while this is acceptable for spike-counting, comparison of spike waveforms required an estimate of the underlying fractional change in fluorescence, ΔF/F. To the extent that resting potentials were approximately the same between cells, ΔF/F provided a measure of relative changes in voltage and thus enabled comparisons of spike amplitude and after-hyperpolarization.

A naïve algorithm would simply fit the clean but uncalibrated ICA-generated trace to a noisy trace with true ΔF/F generated from averaging the raw signal over the entire area of the cell. That is, we might take the average trace 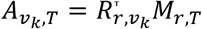 for a given cell and scale the uncalibrated trace to match: 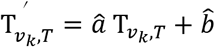 where 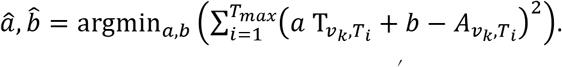 Then the baseline (the F in ΔF/F) would come from low pass filtering this new trace 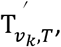 and spikes (the ΔF) would be measured relative to this baseline. The problem is that the average trace 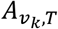 may contain crosstalk from other cells. We therefore sought a better way to estimate an unbiased offset and amplitude for the fluorescence signal.

Instead of using the entire cell mask (𝑅_*r,v*_) to calculate 𝐴_*v,t*_, we identified spatial regions which were unobstructed by neighboring cells. For a cell indexed by 𝑣_*k*_ we calculated the “pure” mask by excluding all the pixels at which other cells were present:

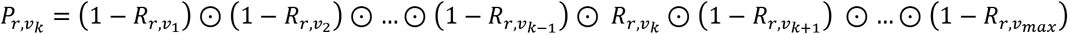

where ⊙ represents elementwise multiplication. In certain rare cases a cell had no regions which were not shared by others. These cells were excluded. To further refine our mask, we identified those pixels within the pure region which matched the ICA-derived trace as closely as possible. We first high pass filtered the movie and the ICA-derived trace in the time domain with a mean-subtracted Gaussian filter set to accentuate action potentials (window size of 20 ms, Gaussian with standard deviation of 3 ms), giving the movie 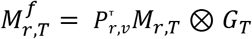 and the trace 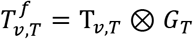 where⊗represents a convolution and 𝐺_*T*_ represents the temporal filter. Following the maximum likelihood pixel weighting algorithm described in Kralj *et al*. (Kralj et al., 2012), we calculated for each pixel of the movie a set of coefficients describing the best fit (by a least squares error) of the ICA-derived trace to the pixel trace. The error on this best fit, normalized the variance of the pixel’s signal, gives an indicator of quality-of-fit: 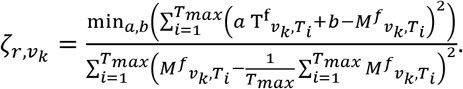 Our final mask was then 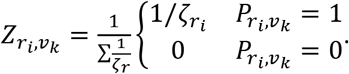 At last, we can construct a new average trace: 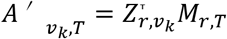 for each cell and perform the fit 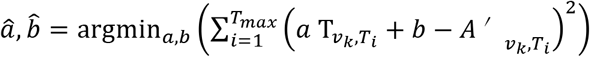 to obtain a trace with real fluorescence units, 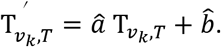

This fluorescence signal still contained artifacts from photobleaching. We estimated the photobleaching baseline using a filter that took the minimum in a sliding window of duration 1 s. Dividing the fluorescence signal by this photobleaching estimate at each time point provided our final ΔF/F trace.

### Estimating morphological parameters

We also used the cell mask, 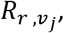 to study the structural morphology of each neuron (see *Morphological analysis* above). Soma and dendrite can be distinguished on the basis of how “thin” they are: pixels on the soma are likely to be surrounded by other on-cell pixels, while pixels on the dendrite are more likely to be adjacent to off-cell pixels. We applied morphological opening to the cell mask 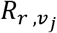 with a disk-shaped structuring element of diameter 5 pixels (∼1 μm). This procedure removed thin areas like dendrites but also shrank the soma. Then we applied morphologically dilation with the same structuring element to restore the soma to its original size without restoring the dendrites. The set of on-cell pixels which are not on the soma, S, gives the dendrites. We can now directly calculate the fraction of the cell that is soma as 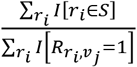 where 𝐼[…] is the indicator function.

### Spike finding

Identification of action potentials in fluorescence traces is complicated by two factors. The first is temporally uncorrelated photon shot noise; the second is low frequency changes in QuasAr2 baseline caused by photobleaching and blue light crosstalk. Action potentials lie at an intermediate frequency between these two noise sources. To deal with low frequency behavior we employed a specialty percentile filter to approximate the baseline fluorescence around each action potential. The filter identified those points in a sliding window (10 frames, 20 ms long) at the 40^th^ percentile and assigned them baseline status. At points not assigned to the baseline, the local baseline value was inferred via linear interpolation. Signal height was then measured relative to this baseline. We found this method to be more robust than conventional linear filtering in regions of rapid baseline change: near stimulus onset, for example, it creates less “delay” in the baseline estimate while still enabling detection of action potentials.

To handle high frequency noise we set a dynamic threshold. The data were median-filtered (with a window of 400 ms) and the difference between the median and the 16^th^ percentile (i.e. −1σ) was taken as a noise estimate. The threshold for spike detection in the high pass filtered data was set to five times this noise estimate. Our conclusions are robust to changes in this threshold multiplier value. Due to high frequency noise, occasionally the same spike crossed the threshold twice. To avoid double-counting, we set a hard limit on the time between two spikes (no less than 18 ms). We found only a very small number of cases where cells approached this frequency limit.

Once spikes were identified, all further characterization was performed on the original (unfiltered) trace.

### Spike parameterization

Spike parameterization is illustrated in Fig. S4 and the approach is justified in Supplemental Discussion. After identifying spikes, we defined windows around each spike on which to perform a functional fit. In the case of an isolated spike, these windows were set to −90 ms and +100 ms from the spike peak. When another spike was within this window, the window was shortened to avoid the neighboring peak. Windows were also truncated to avoid intersection with the stimulus turning on or off.

We fit the pre- and post-peak sections of each action potential separately with a piecewise function with two linear components (termed a “variable hinge”). The function is defined as 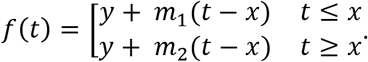 To perform the fit we employed a constrained gradient descent algorithm to minimize least-squares error. Strictly piecewise functions are difficult to optimize, so we approximated *f*(*t*) using logistic functions with fast time constants compared to the steepest action potentials. That is, we used 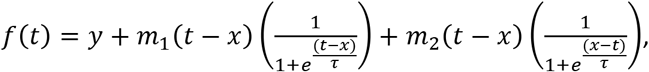, where 𝜏 is the timescale of the logistic (set to *2*×10^−6^ ms). The error function was a Euclidean norm.

After this initial parameterization, we obtained a model for the baseline fluorescence. We first obtained an approximate estimate of the baseline by removing spikes from the fluorescence trace. Points between the pre- and post-peak time constants of an action potential (that is, near the peak) were removed and replaced with a local linear fit to the five points before and after these boundaries. The remaining baseline dynamics at stimulus on and stimulus off were fit separately with variable hinge models, as above.

In the model of the upstroke, the time coordinate of the meeting point was interpreted as the rise time constant and the y coordinate relative to the local baseline was interpreted as the action potential initiation threshold. In model of the downstroke, the time coordinate of the joint was interpreted as the fall time constant and the y coordinate of the joint relative to the local baseline was interpreted as the after-hyperpolarization. We defined action potential height (in units of ΔF/F) as the distance from the peak of each action potential to the local baseline, divided by the baseline. After-hyperpolarization was similarly defined as the distance from the ΔF coordinate of the hinge point to the local baseline, divided by the baseline.

### Statistical methods

In all tests p < 0.01 was considered statistically significant. Nonparametric tests were employed to characterize spontaneous firing rates, the distribution of which showed extreme non-normality. Unpaired t-tests were used to compare firing rates under stimulus, where the average number of action potentials produced was much higher than 0 and the statistics were more closely Gaussian. The Holm-Bonferroni method was used to correct for multiple hypothesis testing where appropriate (in particular, choosing between different mechanisms for high-stimulus differences in firing pattern and identifying significant changes in action potential waveform parameters).

A cell was considered to have entered depolarization block at a particular stimulus intensity if it fired its maximum number of action potentials in the previous stimulus epoch, its maximum number of action potentials was higher than three, and in the current stimulus epoch there were fewer action potentials and more than half of those action potentials occurred in the first half of the stimulus epoch. These constraints were constructed to avoid noise from random small differences in spike timing. Cells which never fired more than three action potentials in a single stimulus were considered constitutively inactive and were excluded from further analysis.

To construct a CheRiff level-insensitive measure of activity, we studied the probability of depolarization block at stimulus *n + 1* conditional on a given number, *Y*, of action potentials at stimulus *n*. We call this probability P(block_n+1_|*Y*_n_). We started with a dataset consisting of all stimulus epochs from all cells which contained more than three action potentials and which occurred prior to either depolarization block or prior to the last (strongest) stimulus. To avoid sampling errors, we further excluded stimulus epochs which contained the maximum number of action potentials (for that cell) but which were not the first stimulus epoch to do so; these would otherwise create a distortion which would depend on whether the last epoch in which the maximum number of action potentials occurred was the last epoch overall.

To calculate P(block_n+1_|*Y*_n_), we employed a generalized linear regression with a binomial distribution and logit linker function, and used cell line (mutant or control), dish recording day (as described in the previous section), the number of action potentials in the current step, and the slope of the F-I curve prior to depolarization block as regression variables. The first two of these variables were treated as categorical. The slope of the F-I curve was included to account for marginal increases in the difference between effective stimulus conductance caused by variations in CheRiff. The given *p* values come from t-tests on these regression coefficients.

To calculate a CheRiff-independent measure of the maximum firing rate, we took only those cells that entered depolarization block to ensure that the maximum firing rate was reached prior to the strongest optogenetic stimulus. We further excluded cells which fired at their maximum rate for more than one stimulus, again to avoid sampling error. We then fit a linear regression to the maximum firing rates with cell line and dish recording day as categorical (non-interacting) regression variables. The given *p* values come from t-tests on these regression coefficients.

In our study of differences in action potential waveform, we first averaged together waveform parameters for all action potentials which occurred later than 100 ms after blue light onset within each stimulus epoch (during this initial period the rising baseline caused significant distortion). Our dataset then consisted of waveform parameters from each stimulus epoch that occurred at or before the first maximum in the F/I curve. We employed a linear regression with coefficients for cell line (mutant or control), dish recording day, the number of action potentials in the current step, and the stimulus epoch number (to account for residual distortions caused by blue light cross-talk). The first two variables were treated as categorical. The given *p* values come from t-tests on these regression coefficients. All statistical analysis was done in MATLAB.

### Simulations

We evaluated a Hodgkin-Huxley type model with parameters taken from Powers and co-workers’ simulations of human motor neurons (Powers and Heckman, 2015, Powers et al., 2012). (The parameters described in the publications (Powers and Heckman, 2015, Powers et al., 2012) are incorrect and do not give spiking behavior; we instead used the parameters in the associated NEURON simulation files provided in ModelDB). The sodium channel activation parameters were adjusted to reflect the recording temperature in our protocol and the slow inactivation component was dropped for simplicity. 𝐾_*v*_-activation parameters in Powers *et al*. were taken from a study (Kuo et al., 2006) in which recordings were performed at the same temperature as in our experiments, and so were not adjusted. Basal K_*v*_*7* conductances were doubled from 1 mS/cm^2^ to *2* mS/cm^2^.

We used the Hodgkin-Huxley equation

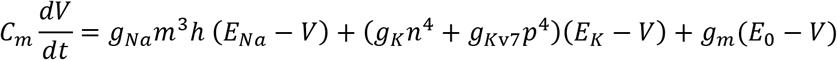

Here 𝑔_*m*_ is a combination of basal leak conductance and CheRiff input conductance. The inactivation particles’ time evolution followed the form

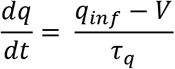

with the parameters

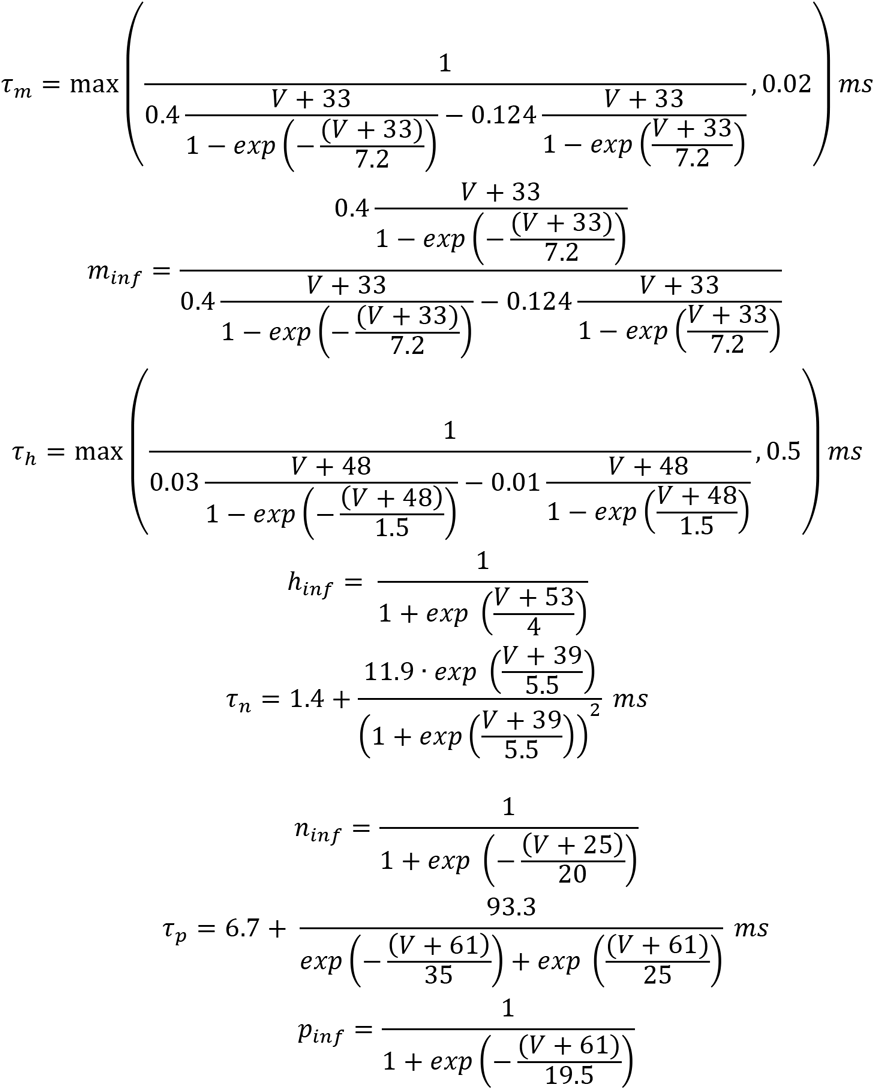

where 𝑉 is in millivolts. The ionic reversal potentials were set to

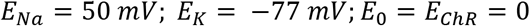

where 𝐸_0_ is the leak reversal potential and 𝐸_*chR*_ is the channelrhodopsin (CheRiff) reversal potential. Our model was evaluated in MATLAB using ode15s to handle stiff behavior, with a relative tolerance of 0.003.

Hodgkin-Huxley-type equations model a differential patch of membrane. We sought to account for the effects of variations in cell size and morphology without creating a full compartment or cable model. Since the active channels involved in action potential production are localized, an increase in cell size can produce an increase in total capacitance without a proportional increase in total active channel conductance. We therefore allowed the lumped capacitance 𝐶_*m*_ to be a tunable parameter and set it to produce maximum firing rates in the range of those seen experimentally (a maximum firing rate before depolarization block of 10-20 action potentials within a 500 ms stimulus); in the simulations shown, it is 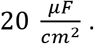 This represents an effective capacitive load, and is not intended to represent capacitance per geometrical membrane area.

For each set of conductance parameters, we allowed the system to evolve to steady state for 10 seconds prior to delivering stimuli. To identify spikes in the simulation, we looked for peaks 25 mV above the steady-state baseline (obtained via a specialty ordinal filter, as discussed in *Spike finding*). Depolarization block was defined in the same way as in the real data. For step stimulations, the input conductance 𝑔_*m*_ was turned on with a time constant of 1 ms. When the cell reached steady state (without entering depolarization block) we obtained more precise measurements of firing rate by running the simulation for 2500 ms and then multiplying the number of action potentials found by 1/5 to obtain the expected number of action potentials per 500 ms, as displayed in Fig. 6.

Spike waveforms were parameterized using the same algorithm as was applied to real data. Simulated voltage time traces were down-sampled and their action potentials parameterized using the same models as applied to the real data. We then fit a linear regression on the average waveform parameter from each step with control variables for the channel conductance and the input conductance (Table S3).

Bean, B.P. (2007). The action potential in mammalian central neurons. Nature Reviews Neuroscience *8*, 451-465.

